# Unique fingerprint of marine ectotherm body size change during hyperthermal crises

**DOI:** 10.1101/2025.07.25.666730

**Authors:** Paulina S. Nätscher, Kenneth De Baets, Wolfgang Kiessling

**Author notes:** Paulina S. Nätscher **Email:**. Shared senior authorship (Kenneth De Baets, Wolfgang Kiessling). **Author Contributions:** PSN, KDB and WK conducted the literature search and collected the data. All authors also designed and conducted the analyses, and were involved in writing and editing the manuscript. **Competing Interest Statement:** The authors declare no competing interests.

## Abstract

The term ‘Lilliput Effect’ describes a substantial decrease in the average body size of fossil assemblages during major environmental perturbations in Earth’s history, which is reported in many paleontological studies. The limited regional, temporal and taxonomic focus of most studies, however, has sparked discussions concerning its generality. Additionally, even though a negative relationship between warming and body size has been established in recent marine ectotherms, the environmental and mechanistic drivers of the Lilliput effect are still debated.

We compiled close to 9000 body size changes from fossil, historical and modern body size studies, to show that a decrease in body size is indeed a general response of marine ectotherms to environmental crises. The magnitude and temporal variability of size changes at the species-level are significantly higher during hyperthermal than non-hyperthermal events, suggesting differing mechanisms of body size decrease depending on the environmental stressor. Our results further show that ancient environmental perturbations with a higher magnitude of warming were associated with a greater dwarfing. This implies that warming was a major driver of body size decreases during hyperthermal events throughout the Phanerozoic, and future warming will impact current trajectories of body size reduction in modern marine ectotherms.

**Significance Statement:** Dwarfing is a common response of marine ectotherms to both anthropogenic warming and similar environmental crises throughout Earth’s history. However, the underlying mechanisms and parallels between fossil and modern observations are still debated. We show, using newly compiled data from fossil, historical and modern studies, that warming events elicit especially strong species body size changes, suggesting similar mechanisms linking modern and fossil body size reductions to warming stress. A correlation between the magnitude of warming during ancient crises and the observed dwarfing indicates that trajectories of body size reduction can be projected into the future depending on climate-change scenarios.

## Introduction

Many studies of body size in the fossil record report that marine organisms have decreased in mean body size during environmental perturbations (1–4). This pattern is widely referred to as the Lilliput effect (5), and is observed across several taxa and past environmental disturbances, but not universally so (6–9). As the vast majority of Lilliput studies focus on single taxa or single localities on relatively short time frames, we still lack a clear understanding on the generality and magnitude of the Lilliput effect across global environmental perturbations.

Instances of the Lilliput effect have been reported in the context of most substantial environmental and evolutionary crises, regardless of differences in the crises’ proximate causes. Potential environmental drivers for the Lilliput effect have been heavily discussed, with most paleontological studies identifying decreased productivity and deoxygenation as the most likely drivers for body size decreases in fossil assemblages (1, 10). In contrast, biological experiments and observations consistently document a negative relationship between marine ectotherm adult body size and seawater temperature, which is referred to as the temperature-size rule (11–13).

Several evolutionary crises throughout the Phanerozoic were probably ultimately caused by Large Igneous Provinces, which evoked global warming through greenhouse-gas emissions similar to the current anthropogenic climate crisis (14–17). Such ancient hyperthermal events provide an opportunity to compare past and modern impacts of climate change on body size changes, especially with regards to differences in time-scales and taxonomic scope (12, 18–21).

We compiled a dataset of nearly 9000 body size change records, from over 1.6 million specimen measurements from paleontological, historical and modern data, to test two hypotheses relating to the generality and mechanisms underlying the Lilliput effect, as well as the role of environmental stressors like warming in driving body size responses:

1. A decrease in body size in fossil assemblages (Lilliput effect) is a general pattern observed in marine ectotherms during most environmental or evolutionary crises throughout the Phanerozoic.
2. The way climate-related stressors have impacted the growth and body size of marine invertebrates in the fossil record, is similar to what we know from modern experimental studies. Based on the temperature-size rule observed in modern marine ectotherms, we hypothesize that crises associated with substantial hyperthermal stress should elicit a stronger body size response within marine ectotherm species than crises that are not driven by warming.

## Results

### Crisis Phases and individual events

We used log response ratios (lnRR) to describe body size changes trough time from time series reported in 171 publications (SI Appendix I, II). We investigated body size changes across the Phanerozoic including background times and ten major environmental and evolutionary crises (Fig. 1, Table 1). Crises include the Big Five mass extinctions and hyperthermal crises (22, 23), around which our dataset is clustered (Fig. S1, Table 1) (24, 25). The overall median body size change in crisis intervals was significantly negative (Fig. 1, Table 1), indicating a consistent presence of dwarfing across the studied crisis intervals. This suggests a general pattern that is consistent with the Lilliput effect in the broad sense (1). For clarity, we hereafter use the term “dwarfing” to refer to a negative shift in median body size, from within-species changes to shifts at the assemblage level. The median body size change during background and rebound phases (45), which we refer to as recovery in this study, was significantly positive (Table 1). The body size responses differed only slightly among ten environmental crises over the last 500 Myr (Fig. 1, Table 1). The median size change in background intervals just before the crises or after the recovery was slightly positive for almost all investigated events, except for the OAE2 (Table 1). Furthermore, the median body size changes during crises were always negative, except for the OAE2, and were followed by positive body size shifts in recovery times, except for the Late Devonian (Fig. 1, Table 1). The latter could be explained by the multiple extinction pulses that make up the Late Devonian mass extinction, or by the sustained origination of predominantly small species, observed in vertebrates (46). The expected pattern of significant dwarfing during the crisis, and a significant body size increase in their aftermath, was observed in the end-Permian and end-Triassic mass extinctions, the PETM, and the historical and modern observations (Table 1). The body size declines during the crisis interval, followed by an increase, albeit not significant, in the recovery were also observed during the Late Ordovician mass extinction, the Pliensbachian-Toarcian, the OAE1 and the K-Pg (Fig. 1, Table 1). The observed patterns in the individual crises were robust to a jackknife cross-validation, except for one publication (28), without which the median body size response to the end-Permian would have been significantly more negative. This publication observed a body size decline in a prolonged crisis interval leading up to the extinction event. A very similar pattern, with dwarfing during the crisis and body size increase in the recovery phase, was observed across most of the abundant taxonomic groups in our dataset (Fig. S2).

**Table 1.** Median body size changes (log response ratios (lnRR)) during background, crisis and recovery times of different crises as shown in Fig. 1. One-sample Wilcoxon signed rank test results show medians significantly (p < 0.05) below (-) or above (+) zero; bold values: p < 0.01).

|  | lnRR Background | lnRR Crisis | lnRR Recovery |
| --- | --- | --- | --- |
| A) ALL | **0.003 (+)** | **-0.038 (-)** | **0.071 (+)** |
| B) Late Ordovician (e.g., 26, 27) | 0.033 (+) | -0.182 | 0.042 |
| C) Late Devonian (e.g., 6, 7) | 0.033 | -0.027 | -0.177 (-) |
| D) End-Permian (e.g., 28–30) | 0.004 | **-0.080 (-)** | 0.288 (+) |
| E) End-Triassic (e.g., 31, 32) | 0.071 | -0.148 (-) | 0.179 (+) |
| F) Pliensbachian-Toarcian (e.g., 33–35) | 0.003 | -0.055 | **0.089 (+)** |
| G) OAE1 (e.g., 36) | 0.005 | -0.033 | 0.279 |
| H) OAE2 (e.g., 37, 38) | -0.001 | 0.003 | 0.015 |
| I) K-Pg (e.g., 39, 40) | 0.004 | **-0.096 (-)** | 0.012 |
| K) PETM (e.g., 3, 41) | 0.003 | -0.161 (-) | 0.014 (+) |
| L) Historical & Modern (e.g., 42–44) | 0.011 | **-0.024 (-)** | 0.201 (+) |

**Figure 1.**
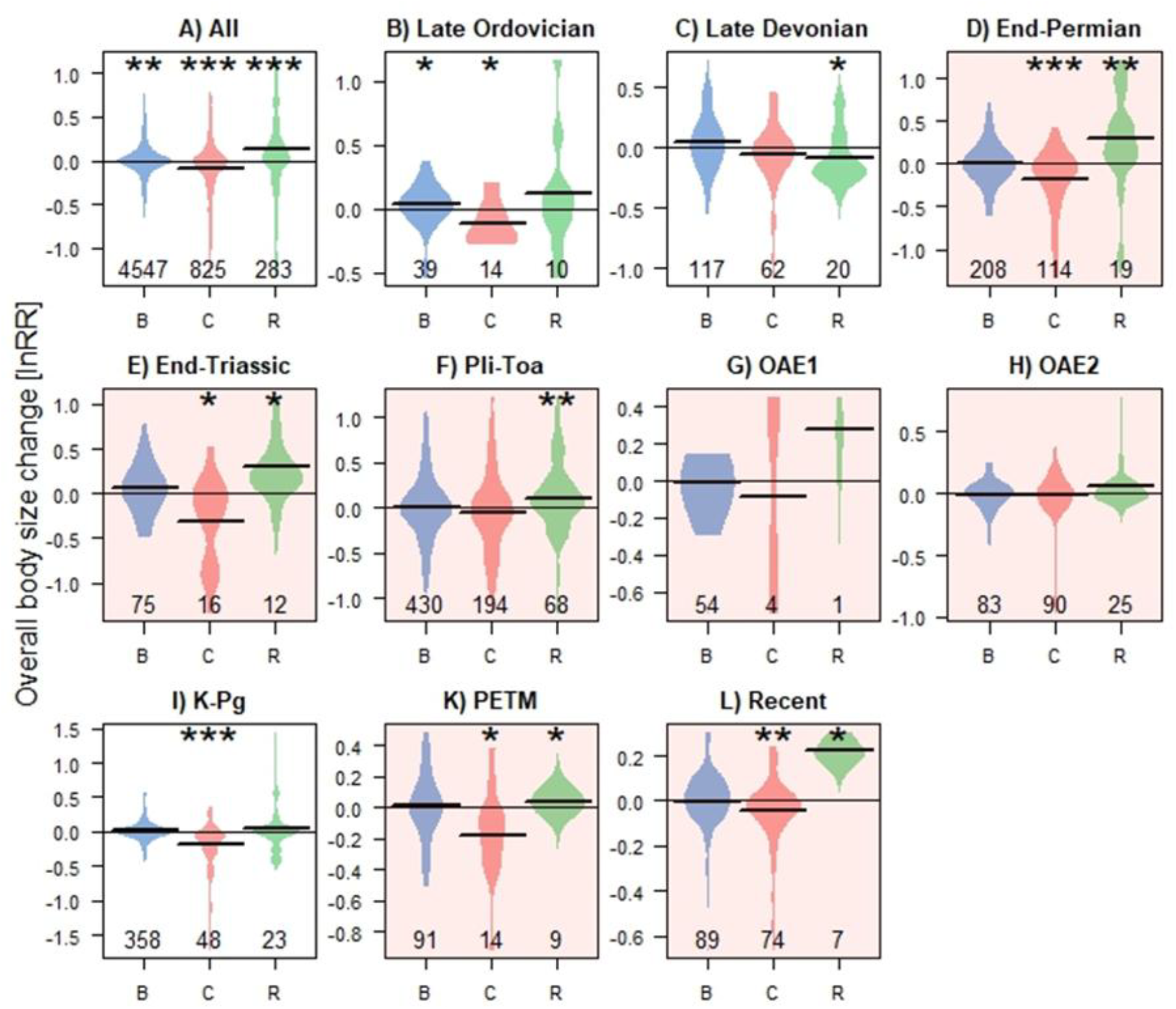
Distribution of overall body size changes (lnRR) during background (blue), crisis (red), and recovery (green) intervals across different Phanerozoic events with outliers removed. Beanplot A) shows all datapoints, while B) to L) show individual events (B: Late Ordovician, C: Late Devonian, D: End-Permian, E: End-Triassic, F: Pliensbachian-Toarcian, G: OAE1, I: OAE2, K: K-Pg, L: Recent). Hyperthermal events (D, E, F, G, H, K, L) are indicated by an orange background. Asterisks indicate medians that differ significantly from zero (^*^ *p* < 0.05, ^**^ *p* < 0.01, ^***^ *p* < 0.001).

### Temporal and taxonomic scaling

There were no significant differences in the overall body size response among five different taxonomic levels (species, genus, family, order, class, assemblage), nor six temporal scales (years, centuries, bed, zone, stage, epoch) that we considered in this study (Tables S1, S2). However, the magnitude of the absolute body size changes rose with increasing taxonomic as well as temporal scale (Fig. S3, Tables S3, S4), indicating the presence of a slight scaling effect (47). This increase in the absolute effect size on broader scales was only broken on the largest taxonomic level, with observations from the class to community scale, where the response magnitude decreases again. The observed pattern of body size response magnitudes across temporal scales was robust, when only species level responses were considered (Tables S2, S4).

### Difference between hyperthermal and non-hyperthermal crises

The Lilliput effect was a uniform response during crises, regardless of whether a crisis was driven by warming or not (Fig. S4). However, when contrasting further characteristics of body size responses of hyperthermal crises and non-hyperthermal crises, a distinct signal of hyperthermal crises emerged. The magnitude (absolute body size change) and volatility (the temporal variability) of body size responses were much larger for hyperthermal crises (Fig. 2). The median magnitude of body size changes during hyperthermal crises was twice that during non-hyperthermal crises (abs(lnRR_hyp_) = 0.144, abs(lnRR_non-hyp_) = 0.065, Wilcox *p* < 0.0001, Fig. 2). Similarly, the median volatility of body size responses during hyperthermal crises was more than twice as high as during non-hyperthermal crises (volatility_hyp_ = 0.274, volatility_non-hyp_ = 0.104, Wilcox *p* = 0.0004). In both cases of magnitude and volatility, the difference in effect size between hyperthermally and non-hyperthermally driven crises became even more pronounced, when only body size changes at the species level were considered (Fig. 2). While the magnitude and volatility of within-species and above-species responses are comparable during hyperthermal events, non-hyperthermal events exhibit significantly stronger and more volatile body size changes above the species-level than within species (magnitude Wilcox *p* < 0.0001, volatility Wilcox *p* = 0.019) (Fig. 2). Within hyperthermal events, a higher magnitude of warming correlated significantly with a higher degree of dwarfing (Fig. 3), suggesting that more negative body size changes occur during events with more pronounced warming.

**Figure 2.**
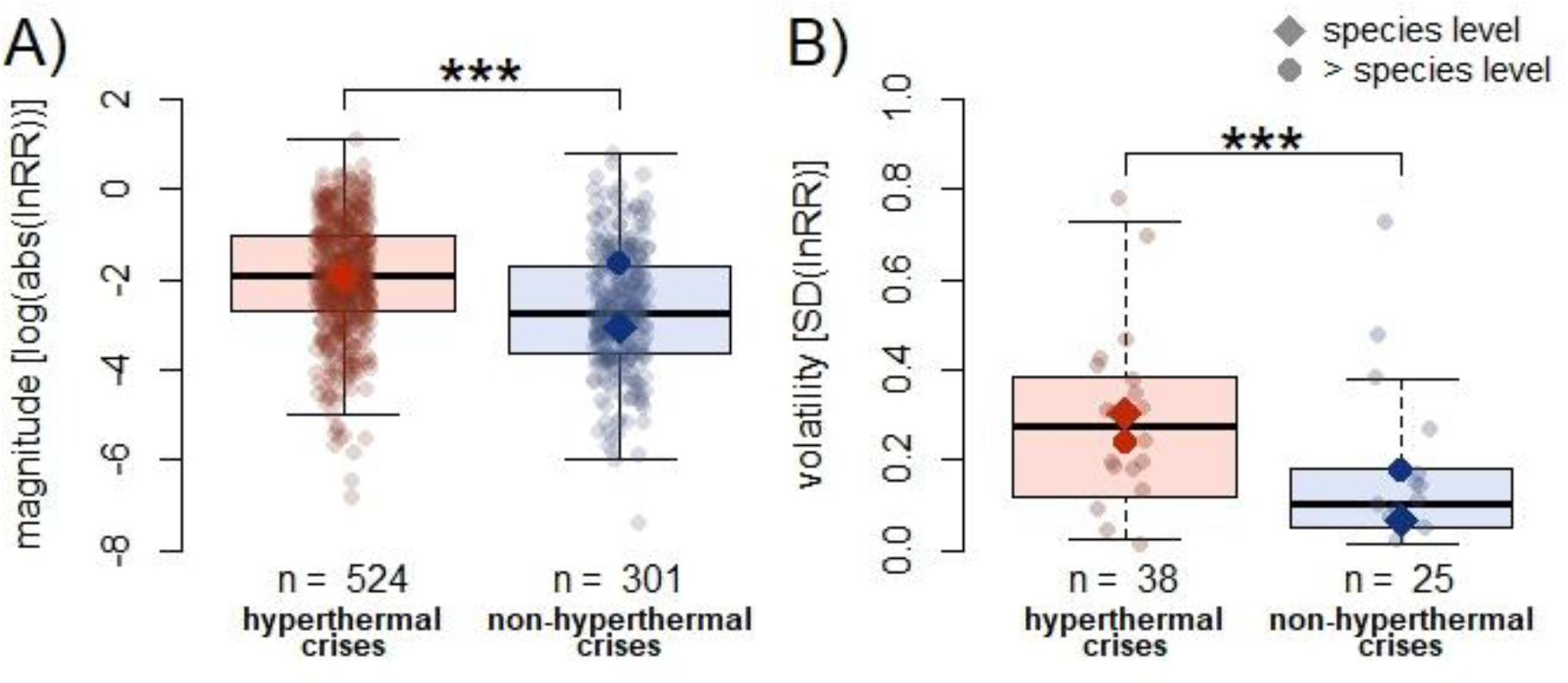
Boxplots of the A) magnitude and B) volatility of body size changes (lnRR) during hyperthermal (red) and non-hyperthermal (blue) crisis intervals depicted in A) absolute log ratios (log-transformed for easier visual distinction) and B) the standard deviation (SD) of crisis body size changes within a time series. Asterisks indicate highly significant (p < 0.0001) statistical differences between the medians in hyperthermal and non-hyperthermal events. Squares and circles show medians of the within-species and above-species level, respectively.

**Figure 3.**
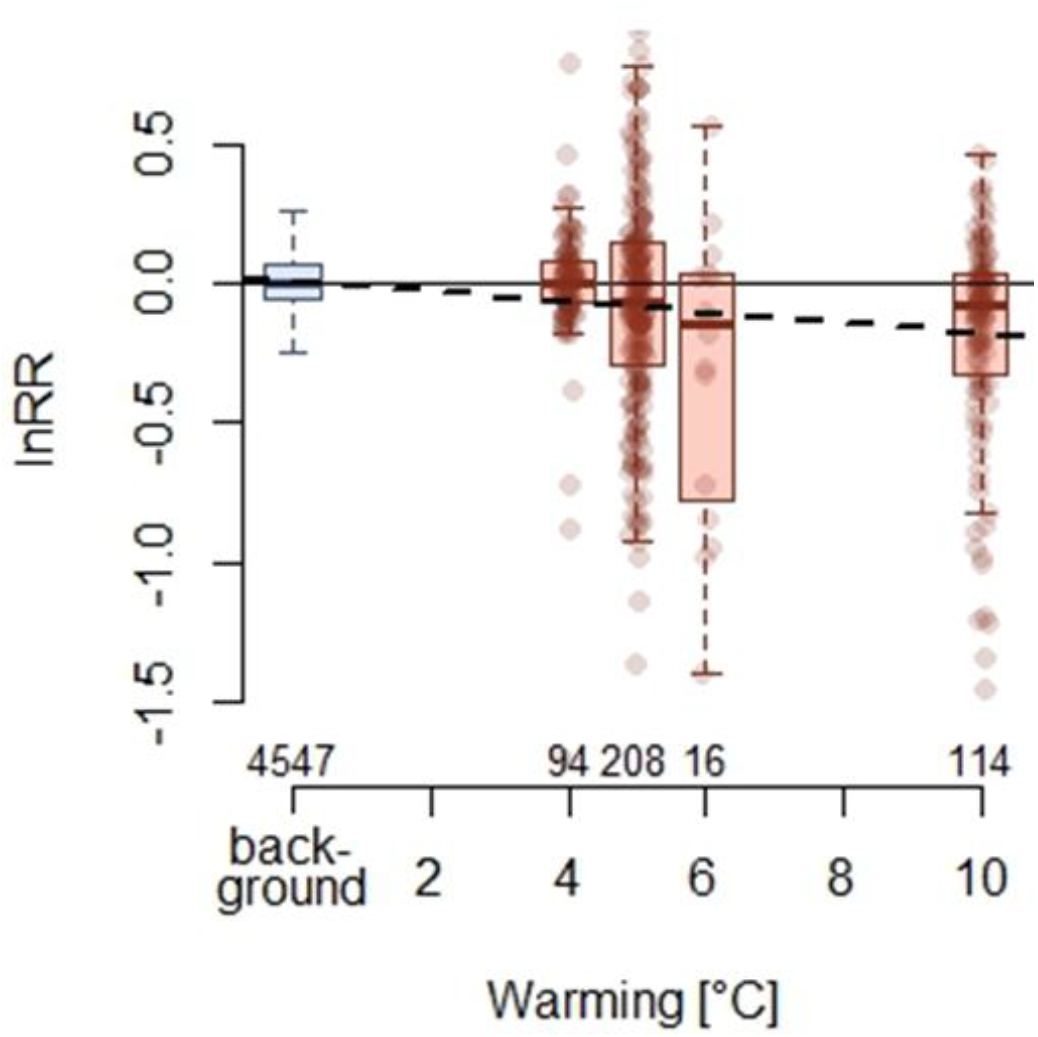
Boxplot of body size changes (lnRR) with degrees (°C) of warming (23), individual data points of hyperthermal body size changes are plotted and sample sizes shown underneath the boxplots. A regression between log response ratios (lnRR) and degrees of warming (°C) shows a modest but significant relationship between body size decline and magnitude of warming (slope = -0.018, R^2^ = 0.018, *p* < 0.0001). A boxplot of all background body size changes is shown in blue on the left.

## Discussion

### Body size decreases are a general response of marine ectotherms in crisis events

We show that dwarfing is a general response of most marine ectotherm groups to most environmental or evolutionary crises throughout the Phanerozoic, regardless of whether they were associated with warming or not (Fig. 1). Crisis-induced declines in body size are usually followed by an increase during the recovery interval (Fig. 1). The most notable exception to this is the Late Devonian mass extinction, after which body sizes continued to decrease. This pattern was observed before and found to be due to a long-term radiation of predominantly small taxa (46). Reductions of body size are reported at all taxonomic and temporal scales ranging from within-species to across classes and from days to geological epochs. Different mechanisms likely operate at different scales, ranging from physiological and ecological to evolutionary (21, 34, 48, 49), however pin-pointing the exact mechanism underlying the body size changes is difficult. We assume that ecological and physiological processes underlie species-level body size changes. Fossil and modern evidence highlights changes in population structure or mechanisms changing the size-at-age, such as ecophenotypic adaptations impacting growth (20, 21, 48, 50). These mechanisms have been shown to lead to substantial body size changes and are reversible, thus resulting in highly volatile changes through time, if the driving stressor fluctuates (21, 50). Body size changes on higher taxonomic levels, such as class or phylum, can be driven by the same mechanisms within abundant species. However, they are also affected by changes in assemblage composition, whether through varying abundances of species or size-biased extinction or origination (20, 51).

Despite our finding of a general Lilliput effect across extinction events, our results can be reconciled with previous studies, which suggested a tendency of smaller genera to disappear during mass extinctions (51–55). Those studies primarily focused on identifying size-biased extinction and origination patterns, i.e., evolutionary body size dynamics of genera. The majority of our dataset, however, focuses on physiological and ecological body size changes over time, with over 60% of the data being recorded at the species level. So, while mass extinctions may be generally selective against smaller genera, we show that especially species decrease in body size across crisis events.

### Hyperthermal stress elicits strong and volatile body size responses in species

Before human activity, rapid global warming was primarily driven by volcanic activity of Large Igneous Provinces (LIPs), which episodically emitted large amounts of greenhouse gases (14–17, 56). Similar to anthropogenic emissions, LIP volcanism caused phases of warming, seawater acidification, ocean stratification, and bottom-water deoxygenation (16, 17, 56, 57), destabilizing marine ecosystems. Warming-induced physiological stress on recent marine ectotherms has been established to be directly related with body size decreases in individuals and assemblages (58–61), and probably had similar effects throughout their evolutionary history (62). Non-hyperthermal events, on the other hand, were associated with stressors like cooling, food scarcity or anoxia (63). Some of these stressors have been shown to affect growth in modern marine organisms only, or especially, when interacting with warming (64, 65). Due to the varied effects of different environmental drivers on organisms and ecosystems, crises that are driven by different stressors may also present different mechanisms underlying body-size decline. This hypothesis is backed by a previous study (66), which found that the established extinction selectivity against smaller taxa during mass extinctions is less pronounced during hyperthermal crises (66). Possibly, because body size decline during hyperthermal events is partially driven by mechanisms unrelated to extinction.

Our results reinforce that body size responses vary among extinction events, depending on the respective environmental driver. Specifically, we show that body size changes, especially those on the species-level, during hyperthermal crises are stronger and more volatile than during non-hyperthermal crises (Fig. 2). We interpret these high volatility species-level body size changes to potentially driven by fast-acting, reversible mechanisms which might occur through temporary phenotypic adaptation to rapid environmental fluctuations (e.g. warming-cooling cycles during volcanic events), which are not present during non-hyperthermal events. This difference between hyperthermal and non-hyperthermal crises cannot be explained by the taxonomic and temporal scaling effects (Fig. S3). To the contrary, non-hyperthermal crises are richer in data on coarse temporal scales, which show higher magnitudes of body size changes than those on finer temporal scales (Fig. S3). If the difference in the magnitude of body size changes between hyperthermal and non-hyperthermal events were driven by scaling effects, non-hyperthermal events should exhibit higher body size change magnitudes. But it is hyperthermal events that show more extreme size responses (Fig. 2).

We also found that the magnitude of warming plays an important role in determining the strength of body size decreases during hyperthermal events, with larger warming magnitudes correlating with greater dwarfing (Fig. 3). The strength of the correlation indicates, however, that other abiotic or biotic factors probably interacted with temperature to mediate organisms’ body size responses. For example, deoxygenation as well as a productivity crises of varying strength would amplify the negative effect of warming (67, 68), while increasing *p*CO_2_ has been shown to mediate negative effects of warming in some cases (69). Along with the lack of significant species body size changes during non-hyperthermal events, we interpret this correlation as a sign that warming and the way it interacts with other stressors have a greater impact on species body size than other stressors alone (Fig. 2). A number of studies reporting body size decreases not in the aftermath of an extinction event, but associated with the gradual warming leading up to hyperthermal crises support this interpretation (28, 70, 71).

Based on our results we suggest two different mechanisms that dominate in driving body size responses during environmental crises. Non-hyperthermal crises, like the Late Ordovician and the K-Pg mass extinctions, likely exerted evolutionary pressure, driving body size changes predominantly on higher taxonomic levels by eliminating or favoring some taxa based on their ability to migrate, or resilience to cooling, fluctuating nutrient conditions, or associated stressors, such as deoxygenation (39). Conversely, the especially strong and volatile species-level body size responses during hyperthermal events suggest the additional presence of underlying ecological or physiological mechanisms, which are fast-acting and reversible. These would not have resulted in the extinction of the species, but in the adaptation of the individuals to the new climatic conditions. During hyperthermal events with extreme warming magnitudes of approximately 10°C, such as the end-Permian mass extinction (23), however, the relatively shallow gradient between dwarfing and the magnitude of warming might indicate that the mechanism underlying body size change switches from species-level phenotypic adaptation high extinction rates, which are biased against smaller genera (Fig. 3) (51, 54, 66). In contrast to previous research focusing on body size driven extinction during mass extinctions, we provide a comparison of background, crisis and recovery patterns in body size changes, with a perspective on surviving species. The more profound impact of warming-driven crises on body size changes might be driven by the interaction of temperature with other stressors, which is established for modern marine ectotherms (61, 72). Our results indicate that the currently observed decline in the body sizes of many marine organisms in response to the rising temperatures will continue, until this adaptive response becomes outweighed by rising extinction rates. This study further highlights the need for a differentiated approach to studying body size changes and diagnosing a Lilliput effect in the fossil record, to make paleontological and biological observations more comparable in the future (34, 48, 49).

## Materials and Methods

### Literature search

In our Web of Science and Google Scholar literature search, we considered publications from the years 1980 to 2024, when titles or abstracts included the keywords “Body Size” or “Size” or “Lilliput” and “Change” or “Trend” or “Variation”. From the resulting papers, we excluded any that focused on freshwater or terrestrial organisms, mammals, insects, reptiles and amphibians. We also excluded studies of body size changes not through time, but in regard to any other variable (e.g., trophic level, location, population density). Body size studies that had been frequently cited in other publications already included in our analysis, were also considered, when they met the criteria. Furthermore, for an article to be included, size measurements had to be available for at least two points in time with a sample size of n ≥ 3 in each sample. Here, we also excluded studies that lacked information about both sample size and an error metric. If either information was given, the paper was included in this study. Overall we collected data from 130 publications that quantify body size changes in marine organisms through time (SI Appendix I). The resulting main table holds 5912 individual changes across a large array of taxonomic groups and ranging from the Cambrian to the Holocene. Our data is clustered around extinction events, such as the end-Permian extinction, the Triassic-Jurassic extinction, the Pliensbachian-Toarcian crisis and the Cretaceous-Paleogene extinction (Fig. S1, Table 1), but measurements from background intervals are also sufficiently present. Body size changes are centered on zero (Fig. S1).

A comprehensive dataset of Hunt et al. (2015), which contains mostly background time series, was used to supplement the background records of body size changes in our study (SI Appendix II) (73). We assessed the 771 time series of morphological data archived in Dryad (https://doi.org/10.5061/dryad.m010p) and extracted those in which the measured trait was either of these: area, log area, length, centroid size or size. Filtering this data for marine environments resulted in 284 individual time series with 2971 individual changes from 41 publications.

### Data acquisition

We compiled body size changes from time series as they are reported in the original publication. Metadata collected directly alongside body size change, included taxonomic information and taxonomic resolution (e.g., phylum, class, genus, species), temporal resolution (from years to geological epochs), system (marine pelagic or shelf). When the same set of body size measurements was presented or otherwise available on varying taxonomic or temporal scales, we entered all available time series, but assigned each resolution step a different natural number as independence indicator. This independence indicator was used to avoid analyzing multiples of data stemming from the same set of specimens in later analyses, by randomly picking and keeping only one taxonomic or temporal resolution from each study. Variables that we collected for both body size measurements within a recorded change were the size measures (mean and standard deviation (SD)) with the original unit, sample sizes, absolute ages and evolutionary phase (background, crisis, recovery). In publications, where the raw data was easily accessible in a supplementary table, we used this data to calculate means and standard deviations (SD) for the body size measurements directly. When only figures were available, we used WebPlotDigitizer-4 to extract means and SDs from the plots (74). In the case that, instead of means and SDs, other forms of statistical descriptors of the data (medians, 95% CI, IQR, se) were given, we used established formulas to transform them into mean and standard deviation (75). An estimate for the absolute age of each body size data point was either taken directly from the publication, when age was supplied, or interpolated between two known age points based on boundary ages from Gradstein *et al*. (2016).

We assigned one of three evolutionary phases (background, crisis, recovery) to each body size measurement based on information on environmental perturbations that was given in each publication. If no environmental or evolutionary crisis, or environmental stress in the case of recent studies, was mentioned in a publication, all measurements were classed as “background”. Body size changes during well-established events of environmental perturbation (incl. the “Big Five” mass extinctions), were automatically classified as “crisis”. The first sample immediately after the perturbations ceased was classed as “recovery”, unless the authors had defined explicit recovery intervals within their recorded time series.

We focused on the Big Five mass extinctions and commonly studied hyperthermal crises, including the current anthropogenic climate crisis (22, 23). Depending on whether or not one of these crises was associated with considerable warming (23), we categorize each event into a hyperthermal or a non-hyperthermal crisis, to assess differences in the body size responses of marine ectotherms based on the presence or absence of hyperthermal stress. Within a temporal bracket of one to two stages around an environmental crisis event (depending on whether the event occurred within a geological stage or coincided with a boundary), all recorded body size changes in which the second measurement is classed as a crisis, were given the categorical descriptor “hyperthermal” or “non-hyperthermal”.

### Analysis

#### Body size changes during ancient environmental crises

Because the types and units of size measurements in the collected data frame vary considerably, we unified units and focused on the log ratios between consecutive measurements to estimate the effect size of body size changes from one interval to the next. We applied the equation

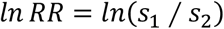

as a metric of size change between two consecutive intervals within each time series. The size values (s_1_, s_2_) are body size measurements in time interval 1 and 2, respectively. Log response ratios (lnRR) are independent of absolute body size and symmetrical for losses and gains. We henceforth refer to this effect size as ‘body size change’. We employed a Kruskal-Wallis test to assess, whether a significant difference exists in the median body size change among the evolutionary phases (background, crisis, recovery) during each individual assessed event as well as the full dataset. A post-hoc Dunn’s test was used to reveal specific pairwise differences between body size changes in background, crisis and recovery phases. We employed one-sample Wilcoxon signed rank tests to assess whether the median body size change in each phase was significantly below or above 0 during each individual assessed event, for each abundant taxonomic group, as well as for the full dataset. We assessed the influence of individual studies on the data by performing a jackknife cross-validation, where the same statistics were calculated for subsets, from which one study was iteratively removed (77). For all analyses, we used a significance threshold of α = 0.05, and *p*-values were corrected to account for a false discovery rate (78).

#### Differences between hyperthermal and non-hyperthermal crises

We assessed differences in the body size responses of marine ectotherms between hyperthermally induced crises and those that are not associated with warming. Here we considered true change, magnitude of change (absolute effect size), as well as volatility of body size changes within a time series as descriptors of body size patterns accompanying environmental or evolutionary crises. To assess the volatility of body size changes we calculated the standard deviation of the effect sizes recorded from consecutive crisis points of each individual independent time series (79). Volatility was calculated using the following formula

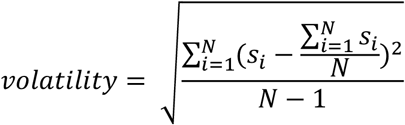

where N is the number of crisis points in a time series and s an individual crisis body size change. Volatility was only calculated, when a time series offered three or more crisis data points of a single category (hyperthermal vs non-hyperthermal). If a time series was sufficiently long and contained enough crisis points of both hyperthermal and non-hyperthermal events, volatility was calculated for both separately. Wilcoxon tests were used to assess differences in directional body size change, magnitude and volatility of body size changes between hyperthermal and non-hyperthermal events. We repeated this for subsets of body size changes above and at the species level.

We employed a linear model to test the correlation between the maximum warming magnitude and body size changes. For this, we assigned the maximum warming magnitude of the six hyperthermal events investigated in this study, to all crisis body size changes recorded during these events (23).

#### Temporal and taxonomic scaling

To test whether body size change varies with differing temporal and taxonomic scales, we first categorized each recorded body size change in our dataset into one of five levels of taxonomic resolution (from small to large: species, genus, family, order, class and above) and one of six levels of temporal resolution (from small to large: years, centuries, bed/sample, formation/zone, stage, epoch). We employed a Kruskal-Wallis test to test for significant differences among the groups and assessed concrete differences in median body size change among scaling units using a Dunn’s post-hoc test.

All analyses were conducted in R Studio (v 2024.04.2+764) (80), using R (v 4.4.1) (81). We employed the package dunn.test (v 1.3.6) (82) in analyses, and divDyn (v 0.8.1) (83) and beanplot (v 1.3.1) (84) for graphics.

## Data Availability Statement

All data and code to produce the results from this manuscript are available at https://doi.org/10.5281/zenodo.15096088.

## Acknowledgments

We thank the editor, one anonymous reviewer, as well as Pedro Monarrez for their constructive feedback on an earlier version of this manuscript. We thank Carl Reddin and Elizabeth Dowding for insightful discussions, and Himadri Haldar for entering data. The project was supported by the Deutsche Forschungsgemeinschaft (DFG projects BA5148/1‐2, KI806/15-2) and is embedded in the Research Unit TERSANE (FOR 2332: Temperature-related stressors as a unifying principle in ancient extinctions). KDB was additionally supported by I.3.4 Action of the Excellence Initiative—Research University Programme at the University of Warsaw, funded by the Ministry of Education and Science, Poland.

## References

1. R. J. Twitchett, The Lilliput effect in the aftermath of the end-Permian extinction event. Palaeogeogr. Palaeoclimatol. Palaeoecol. 252, 132–144 (2007).

2. J.L. Martínez-Díaz, et al., Lilliput effect in a retroplumid crab (Crustacea: Decapoda) across the K/Pg boundary. J. South Am. Earth Sci. 69, 11–24 (2016).

3. T. Yamaguchi, R. D. Norris, A. Bornemann, Dwarfing of ostracodes during the Paleocene-Eocene Thermal Maximum at DSDP Site 401 (Bay of Biscay, North Atlantic) and its implication for changes in organic carbon cycle in deep-sea benthic ecosystem. Palaeogeogr. Palaeoclimatol. Palaeoecol. 346, 130–144 (2012).

4. P. Rita, P. Nätscher, L. V. Duarte, R. Weis, K. De Baets, Mechanisms and drivers of belemnite body-size dynamics across the Pliensbachian-Toarcian crisis. R. Soc. Open Sci. 6, 190494 (2019).

5. A. Urbanek, Biotic crises in the history of upper silurian graptoloids: A palaeobiological model. Hist. Biol. 7, 29–50 (1993).

6. K. R. Brom, M. A. Salamon, P. Gorzelak, Body-size increase in crinoids following the end-Devonian mass extinction. Sci. Rep. 8, 1–7 (2018).

7. C. Girard, S. Renaud, Disentangling allometry and response to Kellwasser anoxic events in the Late Devonian conodont genus Ancyrodella. Lethaia 41, 383–394 (2008).

8. M. Leu, H. Bucher, N. Goudemand, Clade-dependent size response of conodonts to environmental changes during the late Smithian extinction. Earth-Science Rev. 195, 52–67 (2019).

9. P. S. Nätscher, J. Gliwa, K. De Baets, A. Ghaderi, D. Korn, Exceptions to the temperature–size rule: no Lilliput Effect in end-Permian ostracods (Crustacea) from Aras Valley (northwest Iran). Palaeontology 66, 1–16 (2023).

10. P. J. Harries, P. O. Knorr, What does the “Lilliput Effect” mean? Palaeogeogr. Palaeoclimatol. Palaeoecol. 284, 4–10 (2009).

11. D. Atkinson, Temperature and organism size: a biological law for organisms? Adv. Ecol. Res. 25, 1–58 (1994).

12. M. Huss, M. Lindmark, P. Jacobson, R. M. van Dorst, A. Gårdmark, Experimental evidence of gradual size-dependent shifts in body size and growth of fish in response to warming. Glob. Chang. Biol. 25, 2285–2295 (2019).

13. A. R. Coghlan, et al., Mean reef fish body size decreases towards warmer waters. Ecol. Lett. 27, 1–12 (2024).

14. T. J. Algeo, J. Shen, Theory and classification of mass extinction causation. Natl. Sci. Rev. 11, nwad237 (2024).

15. M. R. Rampino, K. Caldeira, S. Rodriguez, Sixteen mass extinctions of the past 541 My correlated with 15 pulses of Large Igneous Province (LIP) volcanism and the 4 largest extraterrestrial impacts. Glob. Planet. Change 234, 104369 (2024).

16. R. E. Ernst, N. Youbi, How Large Igneous Provinces affect global climate, sometimes cause mass extinctions, and represent natural markers in the geological record. Palaeogeogr. Palaeoclimatol. Palaeoecol. 478, 30–52 (2017).

17. D. P. G. Bond, P. B. Wignall, Large igneous provinces and mass extinctions: An update. Spec. Pap. Geol. Soc. Am. 505, 29–55 (2014).

18. P. A. Ahti, A. Kuparinen, S. Uusi-heikkilä, Size does matter—the eco-evolutionary effects of changing body size in fish. Environ. Rev. 28, 311–324 (2020).

19. A. R. Baudron, C. L. Needle, A. D. Rijnsdorp, C. Tara Marshall, Warming temperatures and smaller body sizes: Synchronous changes in growth of North Sea fishes. Glob. Chang. Biol. 20, 1023–1031 (2014).

20. M. Daufresne, K. Lengfellner, U. Sommer, Global warming benefits the small in aquatic ecosystems. Proc. Natl. Acad. Sci. U. S. A. 106, 12788–12793 (2009).

21. P. S. Nätscher, G. Dera, C. J. Reddin, P. Rita, K. De Baets, Morphological response accompanying size reduction of belemnites during an Early Jurassic hyperthermal event modulated by life history. Sci. Rep. 11, 14480 (2021).

22. G. L. Foster, P. Hull, D. J. Lunt, J. C. Zachos, Placing our current “hyperthermal” in the context of rapid climate change in our geological past. Philos. Trans. R. Soc. A Math. Phys. Eng. Sci. 376, 20170086 (2018).

23. W. Kiessling, J. A. Smith, N. B. Raja, Improving the relevance of paleontology to climate change policy. Proc. Natl. Acad. Sci. U. S. A. 120, 1–8 (2023).

24. D. M. Raup, J. J. Sepkoski, Mass Extinctions in the Marine Fossil Record. Science 215, 1501–1503 (1982).

25. P. S. Nätscher, K. De Baets, W. Kiessling, Data for Nätscher et al. “Unique fingerprint of marine ectotherm body size change during hyperthermal crises” https:/doi.org/10.5281/zenodo.15096088.

26. B. Huang, D. A. T. Harper, R. Zhan, J. Rong, Can the Lilliput Effect be detected in the brachiopod faunas of South China following the terminal Ordovician mass extinction? Palaeogeogr. Palaeoclimatol. Palaeoecol. 285, 277–286 (2010).

27. M. R. Borths, W. I. Ausich, Ordovician–Silurian Lilliput crinoids during the end-Ordovician biotic crisis. Swiss J. Palaeontol. 130, 7–18 (2011).

28. W. Kiessling, et al., Pre-mass extinction decline of latest Permian ammonoids. Geology 46, 283–286 (2018).

29. Y. Feng, H. Song, D. P. G. Bond, Size variations in foraminifers from the early Permian to the Late Triassic: Implications for the Guadalupian-Lopingian and the Permian-Triassic mass extinctions. Paleobiology 46, 511–532 (2020).

30. Y. Huang, et al., Temporal shell-size variations of bivalves in South China from the Late Permian to the early Middle Triassic. Palaeogeogr. Palaeoclimatol. Palaeoecol. 609, 111307 (2023).

31. L. F. Opazo, R. J. Twitchett, Bivalve body-size distribution through the Late Triassic mass extinction event. Paleobiology 48, 420–445 (2022).

32. J. W. Atkinson, P. B. Wignall, Body size trends and recovery amongst bivalves following the end-Triassic mass extinction. Palaeogeogr. Palaeoclimatol. Palaeoecol. 538, 109453 (2020).

33. S. D. Morten, R. J. Twitchett, Fluctuations in the body size of marine invertebrates through the Pliensbachian-Toarcian extinction event. Palaeogeogr. Palaeoclimatol. Palaeoecol. 284, 29–38 (2009).

34. P. Rita, P. Nätscher, L. V. Duarte, R. Weis, K. De Baets, Mechanisms and drivers of belemnite body-size dynamics across the Pliensbachian–Toarcian crisis. R. Soc. Open Sci. 6, 190494 (2019).

35. V. Piazza, C. V. Ullmann, M. Aberhan, Temperature-related body size change of marine benthic macroinvertebrates across the Early Toarcian Anoxic Event. Sci. Rep. 10, 1–13 (2020).

36. M. A. Rogov, et al., Response of cephalopod communities on abrupt environmental changes during the early Aptian OAE1a in the Middle Russian Sea. Cretac. Res. 96, 227–240 (2019).

37. A. Takahashi, Responses of inoceramid bivalves to environmental disturbances across the Cenomanian/Turonian boundary in the Yezo forearc basin, Hokkaido, Japan. Cretac. Res. 26, 567–580 (2005).

38. K. R. Brom, M. A. Salamon, B. Ferre, T. Brachaniec, K. Szopa, The Lilliput effect in crinoids at the end of the Oceanic Anoxic Event 2: A Case study from Poland. J. Paleontol. 89, 1076–1081 (2015).

39. M. Aberhan, S. Weidemeyer, W. Kiessling, R. A. Scasso, F. A. Medina, Faunal evidence for reduced productivity and uncoordinated recovery in Southern Hemisphere Cretaceous-Paleogene boundary sections. Geology 35, 227–230 (2007).

40. C. E. Sogot, E. M. Harper, P. D. Taylor, The Lilliput effect in colonial organisms: Cheilostome bryozoans at the Cretaceous-Paleogene mass extinction. PLoS One 9, e87048 (2014).

41. D. N. Schmidt, E. Thomas, E. Authier, D. Saunders, A. Ridgwell, Strategies in times of crisis — insights into the benthic foraminiferal record of the Palaeocene – Eocene Thermal Maximum. Phil. Trans. R. Soc. A 376, 20170328 (2018).

42. J. A. D. Fisher, E. C. Rhile, H. Liu, P. S. Petraitis, An intertidal snail shows a dramatic size increase over the past century. Proc. Natl. Acad. Sci. U. S. A. 106, 5209–5212 (2009).

43. R. J. Wilson-Brodie, M. A. MacLean, P. B. Fenberg, Historical shell size reduction of the dogwhelk (Nucella lapillus) across the southern UK. Mar. Biol. 164, 1–9 (2017).

44. R. Elahi, L. P. Miller, S. Y. Litvin, Historical comparisons of body size are sensitive to data availability and ecological context. Ecology 101, 1–10 (2020).

45. D. Jablonski, S. M. Edie, Mass extinctions and their rebounds: A macroevolutionary framework. Paleobiology (2025) 10.1017/pab.2024.13.

46. L. Sallan, A. K. Galimberti, Body-Size reduction in vertebrates following the end-Devonian mass extinction. Science 350, 812–815 (2015).

47. W. Schlager, D. Marsal, P. A. G. van der Geest, A. Sprenger, Sedimentation rates, observation span, and the problem of spurious correlation. Math. Geol. 30, 547–556 (1998).

48. C. P. Abbott, M. Webster, K. D. Angielczyk, Ontogenetic mechanisms of size change: implications for the Lilliput effect and beyond. Paleobiology 50, 130–149 (2024).

49. B. L. Rego, S. C. Wang, D. Altiner, J. L. Payne, Within- and among-genus components of size evolution during mass extinction, recovery, and background intervals: a case study of Late Permian through Late Triassic foraminifera. Paleobiology 38, 627–643 (2012).

50. H. J. T. Hoving, et al., Extreme plasticity in life-history strategy allows a migratory predator (jumbo squid) to cope with a changing climate. Glob. Chang. Biol. 19, 2089– 2103 (2013).

51. P. M. Monarrez, N. A. Heim, J. L. Payne, Mass extinctions alter extinction and origination dynamics with respect to body size. Proc. R. Soc. B Biol. Sci. 288, 1–8 (2021).

52. C. M. Malanoski, A. Farnsworth, D. J. Lunt, P. J. Valdes, E. E. Saupe, Climate change is an important predictor of extinction risk on macroevolutionary timescales. Science 383, 1130–1134 (2024).

53. J. L. Payne, A. M. Bush, N. A. Heim, M. L. Knope, D. J. McCauley, Ecological selectivity of the emerging mass extinction in the oceans. Science 353, 1284–1286 (2016).

54. J. L. Payne, N. A. Heim, Body size, sampling completeness, and extinction risk in the marine fossil record. Paleobiology 46, 23–40 (2020).

55. J. L. Payne, et al., Selectivity of mass extinctions: Patterns, processes, and future directions. Cambridge Prism. Extinction 1, 1–11 (2023).

56. D. P. G. Bond, S. E. Grasby, On the causes of mass extinctions. Palaeogeogr. Palaeoclimatol. Palaeoecol. 478, 3–29 (2017).

57. D. P. G. Bond, Y. Sun, “Global Warming and Mass Extinctions Associated With Large Igneous Province Volcanism” in Large Igneous Provinces: A Driver of Global Environmental and Biotic Changes, 1st Editio, R.E. Ernst, A. Dickson, A. Bekker, Eds. (Wiley & Sons, 2021), pp. 83–102.

58. H. O. Pörtner, C. Bock, F. C. Mark, Oxygen-& capacity-limited thermal tolerance: Bridging ecology & physiology. J. Exp. Biol. 220, 2685–2696 (2017).

59. C. Deutsch, et al., Impact of warming on aquatic body sizes explained by metabolic scaling from microbes to macrofauna. Proc. Natl. Acad. Sci. U. S. A. 119, 1–9 (2022).

60. M. I. Duncan, et al., Oxygen availability and body mass modulate ectotherm responses to ocean warming. Nat. Commun. 14 (2023).

61. W. C. E. P. Verberk, et al., Shrinking body sizes in response to warming: explanations for the temperature–size rule with special emphasis on the role of oxygen. Biol. Rev. 96, 247–268 (2021).

62. C. J. Reddin, P. S. Nätscher, Á. T. Kocsis, H.-O. Pörtner, W. Kiessling, Marine clade sensitivities to climate change conform across timescales. Nat. Clim. Chang. 10, 249– 253 (2020).

63. W. Kiessling, et al., Marine biological responses to abrupt climate change in deep time. Paleobiology (2024) 10.1017/pab.2024.20.

64. J. L. Johansen, et al., Impacts of ocean warming on fish size reductions on the world’s hottest coral reefs. Nat. Commun. 15, 1–17 (2024).

65. J. Ohlberger, T. J. Cline, D. E. Schindler, B. Lewis, Declines in body size of sockeye salmon associated with increased competition in the ocean. Proc. R. Soc. B Biol. Sci. 290, 20222248 (2023).

66. C. J. Reddin, Á. T. Kocsis, M. Aberhan, W. Kiessling, Victims of ancient hyperthermal events herald the fates of marine clades and traits under global warming. Glob. Chang. Biol. 27, 868–878 (2021).

67. J. Bijma, H. O. Pörtner, C. Yesson, A. D. Rogers, Climate change and the oceans - What does the future hold? Mar. Pollut. Bull. (2013) 10.1016/j.marpolbul.2013.07.022.

68. E. Sampaio, et al., Impacts of hypoxic events surpass those of future ocean warming and acidification. Nat. Ecol. Evol. 5, 311–321 (2021).

69. J. Garzke, T. Hansen, S. M. H. Ismar, U. Sommer, Combined effects of ocean warming and acidification on copepod abundance, body size and fatty acid content. PLoS One 11, 1–22 (2016).

70. V. Piazza, L. V. Duarte, J. Renaudie, M. Aberhan, Reductions in body size of benthic macroinvertebrates as a precursor of the early Toarcian (Early Jurassic) extinction event in the Lusitanian Basin, Portugal. Paleobiology 45, 296–316 (2019).

71. Y. Zhang, et al., Significant pre-mass extinction animal body-size changes: Evidences from the Permian-Triassic boundary brachiopod faunas of South China. Palaeogeogr. Palaeoclimatol. Palaeoecol. 448, 85–95 (2016).

72. A. Audzijonyte, et al., Is oxygen limitation in warming waters a valid mechanism to explain decreased body sizes in aquatic ectotherms? Glob. Ecol. Biogeogr. 28, 64–77 (2019).

73. G. Hunt, M. J. Hopkins, S. Lidgard, Simple versus complex models of trait evolution and stasis as a response to environmental change. Proc. Natl. Acad. Sci. U. S. A. 112, 4885–4890 (2015).

74. A. Rohatgi, S. Rehberg, Z. Stanojevic, Webplotdigitizer: Version 4.1 Of Webplotdigitizer (2018) 10.5281/zenodo.1137880.

75. S. P. Hozo, B. Djulbegovic, I. Hozo, Estimating the mean and variance from the median, range, and the size of a sample. BMC Med. Res. Methodol. 5, 1–10 (2005).

76. J. G. Ogg, G. Ogg, F. M. Gradstein, A Concise Geologic Scale, 1st Ed. (Elsevier, 2016).

77. B. Efron, Bootstrap methods: another look at the jackknife. Ann. Stat. 7, 1–26 (1979).

78. Y. Benjamini, Y. Hochberg, Controlling the False Discovery Rate: A Practical and Powerful Approach to Multiple Testing. J. R. Stat. Soc. Ser. B 57, 289–300 (1995).

79. W. Kiessling, Towards an unbiased estimate of fluctuations in reef abundance and volume during the Phanerozoic. Biogeosciences 3, 15–27 (2006).

80. RStudio Team, RStudio: Integrated Development for R (2020).

81. R Core Team, R: A language and environment for statistical computing. R Found. Stat. Comput. (2021).

82. A. Dinno, dunn.test: Dunn’s Test of Multiple Comparisons Using Rank Sums (2024).

83. Á. T. Kocsis, C. J. Reddin, J. Alroy, W. Kiessling, The r package divDyn for quantifying diversity dynamics using fossil sampling data. Methods Ecol. Evol. 10, 735–743 (2019).

84. P. Kampstra, Beanplot: A Boxplot Alternative for Visual Comparison of Distributions. J. Stat. Software, Code Snippets 28, 1–9 (2008).

