## Supplemental Information for "Unique fingerprint of marine ectotherm body size change during hyperthermal crises"

֍ Shared senior authorship

**This PDF file includes:**

Figures S1 to S4
Tables S1 to S4
Caption for Dataset S1
Caption for Software S1
SI References I and II

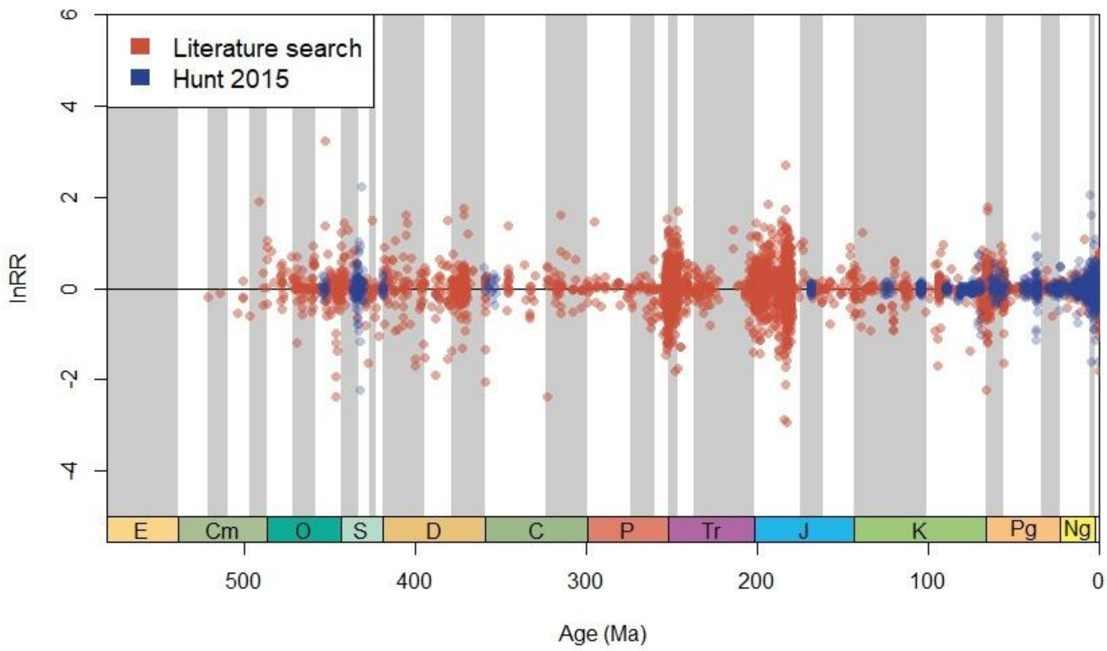


**Fig. S1.** Scatterplot of recorded body size changes, expressed as log response rations (lnRR), throughout the Phanerozoic (red= data from own literature search, blue= Hunt data (56)). The scatter is distributed around 0 and data is densely clustered around some extinction events, especially End-Permian, End-Triassic, Pliensbachian-Toarcian, K-Pg.

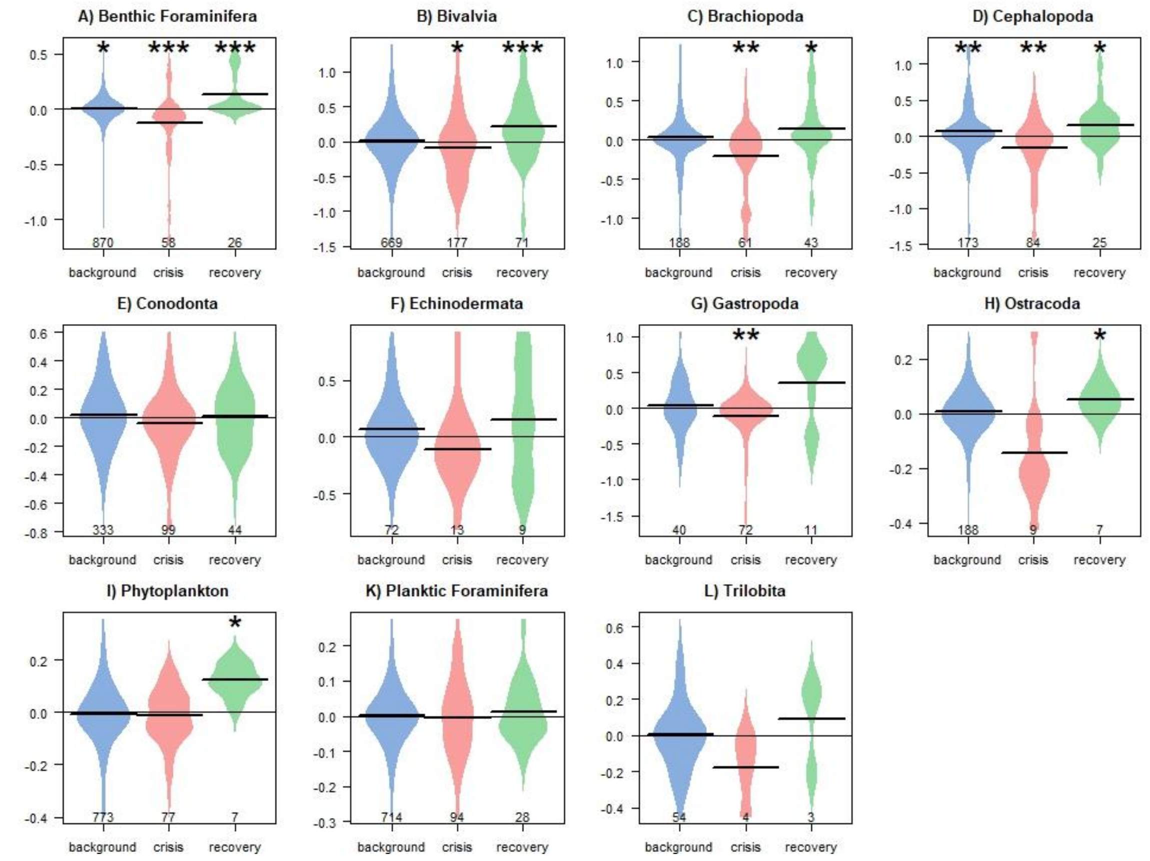


**Fig. S2:** Distribution of overall body size changes (lnRR) of the most abundant taxonomic groups in the dataset during background (blue), crisis (red), and recovery (green) intervals with outliers removed. Beanplots show A) benthic Foraminifera, B) Bivalvia ,C) Brachiopoda, D) Cephalopoda, E) Conodonta, F) Echinodermata, G) Gastropoda, H) Ostracoda, I) Phytoplankton, K) planktonic Foraminifera, L) Trilobita. Asterisks indicate medians that differ significantly from zero (* $p < 0.05$, ** $p < 0.01$, *** $p < 0.001$).

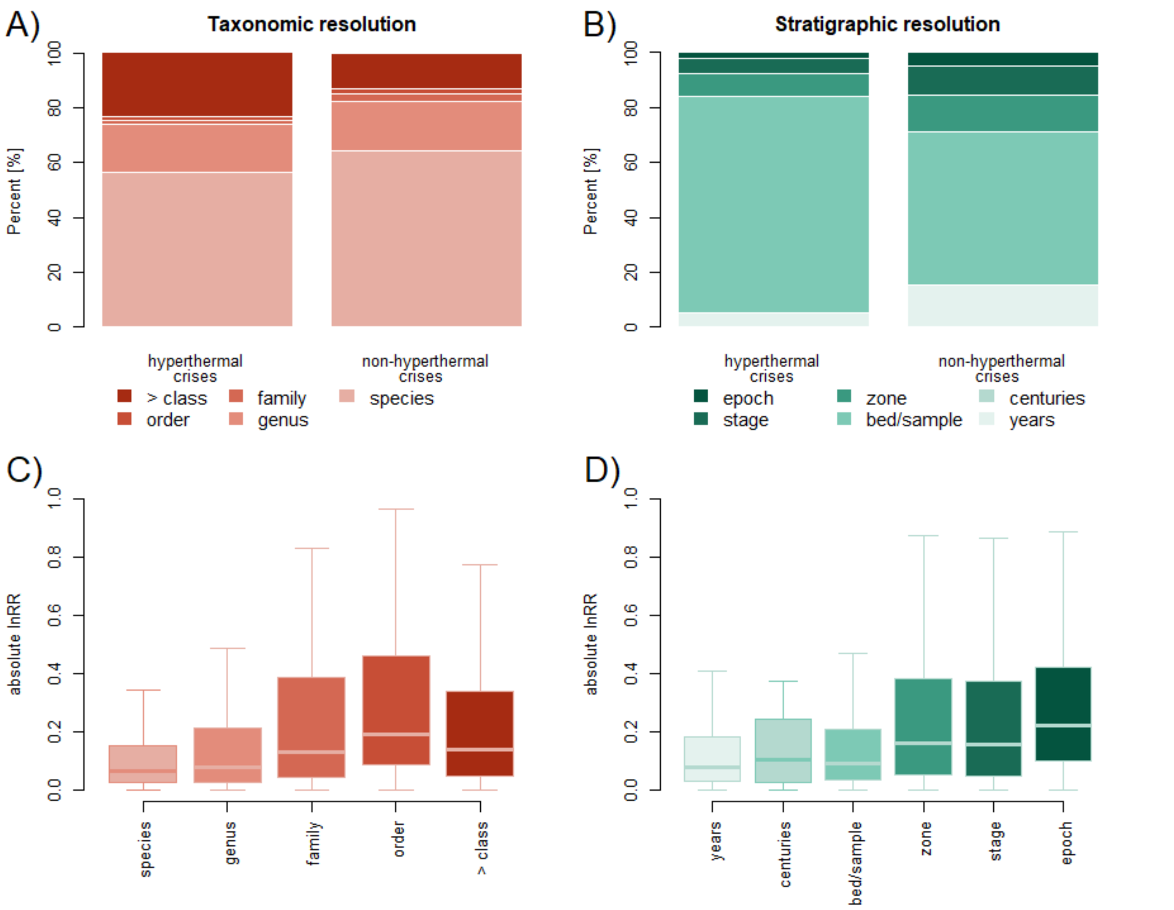


**Fig. S3.** The stacked barplots (A) and (B) show the relative proportions of different taxonomic and stratigraphic scales making up the samples of hyperthermal and non-hyperthermal body size changes. The boxplots (C) and (D) show differences in the magnitude of different body size changes (abs(lnRR)) in taxonomic and stratigraphic scales, respectively.

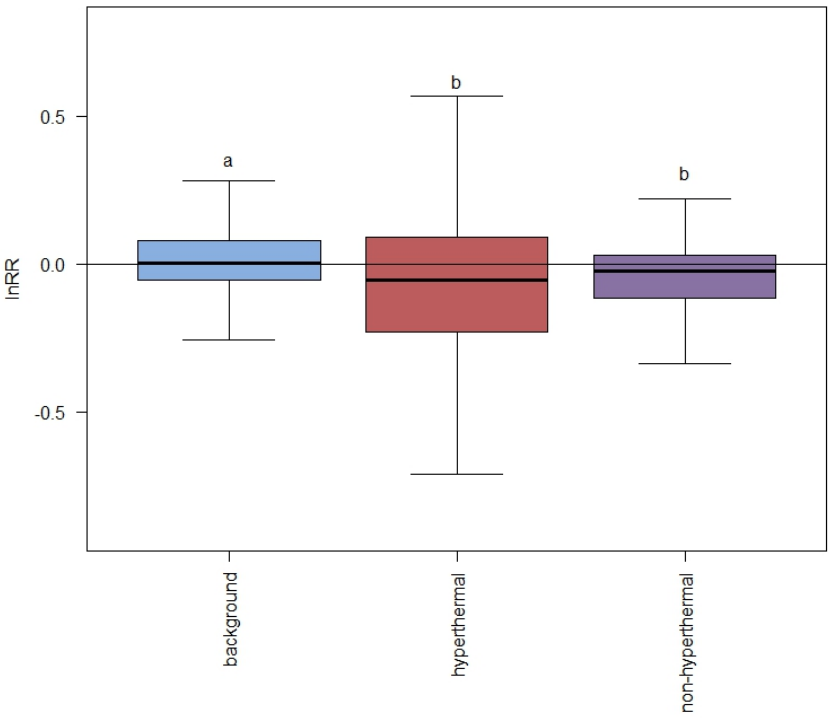


**Fig. S4.** Boxplots showing overall body size changes in background intervals, hyperthermal crises and non-hyperthermal crises. Different letters indicate significant difference (α < 0.05) among groups based on a Kruskal-Wallis test and a post-hoc Dunn's test.

**Table S1**. Results of a post-hoc Dunn's test, comparing differences in body size changes among the five taxonomic levels (species, genus, family, order, class and above), the top cell shows the difference between the row and column mean and the bottom cell contains the pairwise p-value resulting from the Dunn's test. There are no significant differences at α=0.05. Kruskal-Wallis: K-W chi-squared = 6.209, df = 4, *p* = 0.18

| | Species | Genus | Family | Order |
|---|---|---|---|---|
| Genus | 1.292 | | | |
| | 0.245 | | | |
| Family | -0.376 | -0.762 | | |
| | 0.442 | 0.319 | | |
| Order | -1.951 | -2.346 | -0.979 | |
| | 0.128 | 0.095 | 0.273 | |
| Phylum | -0.144 | -1.123 | 0.318 | 1.812 |
| | 0.443 | 0.262 | 0.317 | 0.117 |

**Table S2.** Results of a post-hoc Dunn's test, comparing differences in the overall body size changes among the six temporal scales (years, centuries, bed, zone, stage, epoch), the top cell shows the difference between the row and column mean and the bottom cell contains the pairwise p-value resulting from the Dunn's test (full dataset left side, species subset right side), significant differences (at α=0.05) are indicated by an asterisk. Kruskal-Wallis: all: chi-squared = 5.382, df = 5, p = 0.37, species: chi-squared = 8.600, df = 5, p = 0.13

| | Years | | Centuries | | Bed | | Zone | | Stage | |
|---|---|---|---|---|---|---|---|---|---|---|
| Centuries | -1.063 | -1.157 | | | | | | | | |
| | 0.360 | 0.185 | | | | | | | | |
| Bed | -0.593 | -0.843 | 0.976 | 1.024 | | | | | | |
| | 0.319 | 0.230 | 0.247 | 0.209 | | | | | | |
| Zone | -1.162 | -1.379 | 0.851 | 0.832 | -0.978 | -0.939 | | | | |
| | 0.460 | 0.180 | 0.269 | 0.217 | 0.273 | 0.217 | | | | |
| Stage | -1.173 | -2.218 | 0.840 | 0.229 | -0.979 | -1.980 | -0.060 | -1.405 | | |
| | 0.603 | 0.199 | 0.251 | 0.409 | 0.307 | 0.179 | 0.476 | 0.200 | | |
| Epoch | -1.706 | 1.165 | 0.508 | 1.650 | -1.563 | 1.290 | -1.083 | 1.425 | -1.037 | 1.840 |
| | 0.661 | 0.203 | 0.328 | 0.186 | 0.443 | 0.185 | 0.418 | 0.231 | 0.321 | 0.165 |

**Table S3.** Results of a post-hoc Dunn's test, comparing differences in the **absolute** magnitude of body size changes among the five taxonomic levels (species, genus, family, order, class and above), the top cell shows the difference between the row and column mean and the bottom cell contains the pairwise p-value resulting from the Dunn's test, significant differences (at α=0.05) are indicated by an asterisk. Kruskal-Wallis: K-W chi-squared = 354.337.536, df = 4, $p < 0.0001$

| | Species | Genus | Family | Order |
|---|---|---|---|---|
| Genus | -4.745 | | | |
| | <0.0001* | | | |
| Family | -4.928 | -3.350 | | |
| | <0.0001* | 0.0005* | | |
| Order | -10.205 | -8.128 | -2.824 | |
| | <0.0001* | <0.0001* | 0.003* | |
| Phylum | -16.162 | -9.561 | -0.410 | 3.666 |
| | <0.0001* | <0.0001* | 0.341 | 0.0002* |

**Table S4.** Results of a post-hoc Dunn's test, comparing differences in the absolute magnitude of body size changes among the six temporal scales (years, centuries, bed, zone, stage, epoch), the top cell shows the difference between the row and column mean and the bottom cell contains the pairwise p-value resulting from the Dunn's test (full dataset left side, species subset right side), significant differences (at α=0.05) are indicated by an asterisk. Kruskal-Wallis: all: chi-squared = 212.060, df = 5, p < 0.0001, species: chi-squared = 138.649, df = 5, p < 0.0001)

| | Years | | Centuries | | Bed | | Zone | | Stage | |
|---|---|---|---|---|---|---|---|---|---|---|
| Centuries | -0.131 | -0.070 | | | | | | | | |
| | 0.448 | 0.472 | | | | | | | | |
| Bed | -2.204 | 1.316 | -0.231 | 0.296 | | | | | | |
| | 0.021* | 0.202 | 0.438 | 0.443 | | | | | | |
| Zone | -8.512 | -7.942 | -1.471 | -1.779 | -9.993 | -10.890 | | | | |
| | <0.0001* | <0.0001* | 0.096 | 0.113 | <0.0001* | <0.0001* | | | | |
| Stage | -7.960 | -4.327 | -1.418 | -1.617 | -8.878 | -5.151 | 0.315 | 0.171 | | |
| | <0.0001* | <0.0001* | 0.098 | 0.132 | <0.0001* | <0.0001* | 0.434 | 0.463 | | |
| Epoch | -7.829 | -0.529 | -2.294 | -0.368 | -7.471 | -0.716 | -2.984 | 0.990 | -3.111 | 0.897 |
| | <0.0001* | 0.407 | 0.018* | 0.446* | <0.0001* | 0.355 | 0.003* | 0.302 | 0.002* | 0.308 |

**Dataset S1** (separate file). Dataset of body size measurements and changes extracted from the literature from SI References I and II.

**Software S1** (separate file). R code and R Data file to replicate the analyses and figures presented in this manuscript.

**SI Appendix References I (References resulting from literature search)**


1. Aberhan, M., Weidemeyer, S., Kiessling, W., Scasso, R. A., & Medina, F. A. (2007). Faunal evidence for reduced productivity and uncoordinated recovery in Southern Hemisphere Cretaceous-Paleogene boundary sections. *Geology*, 35(3), 227-230.
2. Anderson, L. C. (2001). Temporal and geographic size trends in Neogene Corbulidae (Bivalvia) of tropical America: using environmental sensitivity to decipher causes of morphologic trends. *Palaeogeography, Palaeoclimatology, Palaeoecology*, 166(1-2), 101-120.
3. Atkinson, J. W., & Wignall, P. B. (2020). Body size trends and recovery amongst bivalves following the end-Triassic mass extinction. *Palaeogeography, Palaeoclimatology, Palaeoecology*, 538, 109453.
4. Atkinson, J. W., Wignall, P. B., Morton, J. D., & Aze, T. (2019). Body size changes in bivalves of the family Limidae in the aftermath of the end-Triassic mass extinction: the Brobdingnag effect. *Palaeontology*, 62(4), 561-582.
5. Barbarin, N., Bonin, A., Mattioli, E., Pucéat, E., Cappetta, H., Gréselle, B., Pittet, B., Vennin, E. and Joachimski, M. (2012). Evidence for a complex Valanginian nannoconid decline in the Vocontian basin (South East France). *Marine Micropaleontology*, 84, 37-53.
6. Baudron, A. R., Needle, C. L., Rijnsdorp, A. D., & Tara Marshall, C. (2014). Warming temperatures and smaller body sizes: synchronous changes in growth of North Sea fishes. *Global change biology*, 20(4), 1023-1031.
7. Becker, R. T. (2005). Ammonoid evolution before, during and after the" Kellwasser-event"—review and preliminary new results. In Global Bio-Events: A Critical Approach Proceedings of the First International Meeting of the IGCP Project 216:“Global Biological Events in Earth History” (pp. 181-188). Berlin, Heidelberg: Springer Berlin Heidelberg.
8. Bell, M.A., Braddy, S.J. (2012). Cope's rule in the Ordovician trilobite family Asaphidae (order Asaphida): patterns across multiple most parsimonious trees. *Historical Biology: An International Journal of Palaeobiology,* 24(3), 223-230
9. Borths, M.R., Ausich, W.I. (2011). Ordovician-Silurian Lilliput crinoids during the end-Ordovician biotic crisis. *Swiss Journal of Palaeontology,* 130(7), 7-18.
10. Brom, K. R., Niedźwiedzki, R., Brachaniec, T., Ferré, B., & Salamon, M. A. (2016). Environmental control on shell size of Middle Triassic bivalve Plagiostoma. *Carnets Geol.*, 16(10), 297.
11. Brom, K. R., Salamon, M. A., Ferré, B., Brachaniec, T., & Szopa, K. (2015). The Lilliput effect in crinoids at the end of the Oceanic Anoxic Event 2: a case study from Poland. *Journal of Paleontology*, 89(6), 1076-1081.
12. Brom, K. R., Salamon, M. A., & Gorzelak, P. (2018). Body-size increase in crinoids following the end-Devonian mass extinction. *Scientific Reports*, 8(1), 9606.
13. Brombacher, A., Elder, L.E., Hull, P.M., Wilson, P.A., Ezard, T.H.G. (2018). Calibration of test diameter and area as proxies for body size in the planktonic foraminifer *Globoconella puncticulata*. *Journal of Foraminiferal Research*, 48(3), 241-245.
14. Caswell, B. A., and Coe, A. L. (2013). Primary productivity controls on opportunistic bivalves during Early Jurassic oceanic deoxygenation: *Geology*, 41(11), 1163-1166.
15. Caswell, B. A., & Dawn, S. J. (2019). Recovery of benthic communities following the Toarcian oceanic anoxic event in the Cleveland Basin, UK. *Palaeogeography, Palaeoclimatology, Palaeoecology*, 521, 114-126.
16. Chen, Y., Neubauer, T. A., Krystyn, L., & Richoz, S. (2016). Allometry in Anisian (Middle Triassic) segminiplanate conodonts and its implications for conodont taxonomy. *Palaeontology*, 59(5), 725-741.
17. Chen, J., Song, H., He, W., Tong, J., Wang, F., and Wu, S. (2018). Size variation of brachiopods from the Late Permian through the Middle Triassic in South China: Evidence for the Lilliput Effect following the Permian-Triassic extinction. *Palaeogeography, Palaeoclimatology, Palaeoecology*, 519, 248-257.
18. Chen, Y., Twitchett, R.J., Jiang, H., Richoz, S., Lai, X., Yan, C., Sun, Y., Liu, X. and Wang, L. (2013). Size variation of conodonts during the Smithian–Spathian (Early Triassic) global warming event. *Geology*, 41(8), 823-826.
19. Daley, G. M. (1999). Environmentally controlled variation in shell size of *Ambonychia* Hall (Mollusca, Bivalvia) in the type Cincinnatian (Upper Ordovician). *Palaios*, 14(6), 520-529.
20. Daufresne, M., Lengfellner, K., & Sommer, U. (2009). Global warming benefits the small in aquatic ecosystems. *Proceedings of the National Academy of Sciences*, 106(31), 12788-12793.

21. Davis, C. V., Myhre, S. E., & Hill, T. M. (2016). Benthic foraminiferal shell weight: Deglacial species-specific responses from the Santa Barbara Basin. *Marine Micropaleontology*, 124, 45-53.
22. Dommergues, J. L., Montuire, S., & Neige, P. (2002). Size patterns through time: the case of the Early Jurassic ammonite radiation. *Paleobiology*, 28(4), 423-434.
23. Elahi, R., Miller, L. P., & Litvin, S. Y. (2020). Historical comparisons of body size are sensitive to data availability and ecological context. *Ecology*, 101(9), e03101.
24. Elahi, R., Sebens, K. P., & De Leo, G. A. (2016). Ocean warming and the demography of declines in coral body size. *Marine Ecology Progress Series*, 560, 147-158.
25. Estes, J. A., Lindberg, D. R., & Wray, C. (2005). Evolution of large body size in abalones (*Haliotis*): patterns and implications. *Paleobiology*, 31(4), 591-606.
26. Falzoni, F., Petrizzo, M. R., & Valagussa, M. (2018). A morphometric methodology to assess planktonic foraminiferal response to environmental perturbations: the case study of Oceanic Anoxic Event 2, Late Cretaceous. *Bollettino della Società Paleontologica Italiana*, 57(2), 103-124.
27. Feng, Y., Song, H., & Bond, D. P. (2020). Size variations in foraminifers from the early Permian to the Late Triassic: implications for the Guadalupian–Lopingian and the Permian–Triassic mass extinctions. *Paleobiology*, 46(4), 511-532.
28. Ferraro, S., Coccioni, R., Sabatino, N., Del Core, M., & Sprovieri, M. (2020). Morphometric response of late Aptian planktonic foraminiferal communities to environmental changes: A case study of *Paraticinella rohri* at Poggio le Guaine (central Italy). *Palaeogeography, Palaeoclimatology, Palaeoecology*, 538, 109384.
29. Fisher, J. A., Rhile, E. C., Liu, H., & Petraitis, P. S. (2009). An intertidal snail shows a dramatic size increase over the past century. *Proceedings of the National Academy of Sciences*, 106(13), 5209-5212.
30. Forel, M. B., Crasquin, S., Chitnarin, A., Angiolini, L., & Gaetani, M. (2015). Precocious sexual dimorphism and the Lilliput effect in Neo-Tethyan Ostracoda (Crustacea) through the Permian–Triassic boundary. *Palaeontology*, 58(3), 409-454.
31. Foster, W.J., Gliwa, J., Lembke, C., Pugh, A.C., Hofmann, R., Tietje, M., Varela, S., Foster, L.C., Korn, D. and Aberhan, M. (2020). Evolutionary and ecophenotypic controls on bivalve body size distributions following the end-Permian mass extinction. *Global and Planetary Change*, 185, 103088.
32. Fuksi, T., Tomašových, A., Gallmetzer, I., Haselmair, A., & Zuschin, M. (2018). 20th century increase in body size of a hypoxia-tolerant bivalve documented by sediment cores from the northern Adriatic Sea (Gulf of Trieste). *Marine Pollution Bulletin*, 135, 361-375.
33. Joral, F. G., Baeza-Carratalá, J. F., & Goy, A. (2018). Changes in brachiopod body size prior to the Early Toarcian (Jurassic) Mass Extinction. *Palaeogeography, Palaeoclimatology, Palaeoecology,* 506, 242-249.
34. Girard, C., & Renaud, S. (1996). Size variation in conodonts in response to the Upper Kellwasser crisis (Upper Devonian of the Montagne Noire France). *CR Acad. Sci.,* 323, 435-442.
35. Girard, C., & Renaud, S. (2008). Disentangling allometry and response to Kellwasser anoxic events in the Late Devonian conodont genus *Ancyrodella*. *Lethaia*, 41(4), 383-394.
36. Grey, M., Finkel, Z. V., Pufahl, P. K., & Reid, L. M. (2012). Evolutionary mode of the ostracod, *Velatomorpha altilis*, from the Joggins Fossil Cliffs UNESCO World Heritage Site. *Lethaia*, 45(4), 615-623.
37. Guinot, G., & Cavin, L. (2018). Body size evolution and habitat colonization across 100 million years (Late Jurassic-Paleocene) of the actinopterygian evolutionary history. *Fish and Fisheries*, 19(4), 577-597.
38. Hall, J. L. O. (2017). Marine bivalve records of Antarctic seasonality and biological responses to environmental change over the Cretaceous-Paleogene mass extinction interval (Doctoral dissertation, University of Leeds).
39. Hallam, A. (1998). Speciation patterns and trends in the fossil record. *Geobios,* 30, 921-930.
40. Hatakeda, K., Suzuki, N., & Matsuoka, A. (2007). Quantitative morphological analyses and evolutionary history of the Middle Jurassic polycystine radiolarian genus *Striatojaponocapsa* Kozur. *Marine Micropaleontology*, 63(1-2), 39-56.
41. He, W., Shi, G. R., Yang, T., Zhang, K., Yue, M., Xiao, Y., ... & Wu, S. (2016). Patterns of brachiopod faunal and body-size changes across the Permian− Triassic boundary: Evidence from the Daoduishan section in Meishan area, South China. *Palaeogeography, Palaeoclimatology, Palaeoecology,* 448, 72-84.

42. Hoving, H. J. T., Gilly, W. F., Markaida, U., Benoit-Bird, K. J., -Brown, Z. W., Daniel, P., ... & Campos, B. (2013). Extreme plasticity in life-history strategy allows a migratory predator (jumbo squid) to cope with a changing climate. *Global change biology*, 19(7), 2089-2103.
43. Huang, B., Harper, D. A., Zhan, R., & Rong, J. (2010). Can the Lilliput Effect be detected in the brachiopod faunas of South China following the terminal Ordovician mass extinction?. Palaeogeography, Palaeoclimatology, Palaeoecology, 285(3-4), 277-286.
44. Huang, Y., Tong, J., Tian, L., Song, H., Chu, D., Miao, X., & Song, T. (2023). Temporal shell-size variations of bivalves in South China from the Late Permian to the early Middle Triassic. *Palaeogeography, Palaeoclimatology, Palaeoecology*, 609, 111307.
45. Huber, R., Meggers, H., Baumann, K. H., Raymo, M. E., & Henrich, R. (2000). Shell size variation of the planktonic foraminifer *Neogloboquadrina pachyderma* sin. in the Norwegian–Greenland Sea during the last 1.3 Myrs: implications for paleoceanographic reconstructions. *Palaeogeography, Palaeoclimatology, Palaeoecology*, 160(3-4), 193-212.
46. Hunt, G., & Roy, K. (2006). Climate change, body size evolution, and Cope's Rule in deep-sea ostracodes. *Proceedings of the National Academy of Sciences*, 103(5), 1347-1352.
47. Hunt, G., Wicaksono, S. A., Brown, J. E., & MacLeod, K. G. (2010). Climate-driven body-size trends in the ostracod fauna of the deep Indian Ocean. *Palaeontology*, 53(6), 1255-1268.
48. Ivany, L. C., Pietsch, C., Handley, J. C., Lockwood, R., Allmon, W. D., & Sessa, J. A. (2018). Little lasting impact of the Paleocene-Eocene Thermal Maximum on shallow marine molluscan faunas. *Science Advances*, 4(9), eaat5528.
49. Jackson, G. D., & Domeier, M. L. (2003). The effects of an extraordinary El Niño/La Niña event on the size and growth of the squid *Loligo opalescens* off Southern California. *Marine Biology*, 142(5), 925-935.
50. Jarrett, M. B. (2016). Lilliput Effect Dynamics across the Cretaceous-Paleogene Mass Extinction: Approaches, Prevalence, and Mechanisms. Graduate Theses and Dissertations
51. Kamikuri, S. (2012). Evolutionary changes in the biometry of the fossil radiolarian *Stichocorys peregrina* lineage in the eastern equatorial and eastern North Pacific. *Marine Micropaleontology* 90-91, 13-28.
52. Kiessling, W., Schobben, M., Ghaderi, A., Hairapetian, V., Leda, L., & Korn, D. (2018). Pre–mass extinction decline of latest Permian ammonoids. *Geology*, 46(3), 283-286.
53. Klug, C., De Baets, K., Kröger, B., Bell, M. A., Korn, D., & Payne, J. L. (2015). Normal giants? Temporal and latitudinal shifts of Palaeozoic marine invertebrate gigantism and global change. *Lethaia*, 48(2), 267-288.
54. Klug, C., Schatz, W., Korn, D., & Reisdorf, A. G. (2005). Morphological fluctuations of ammonoid assemblages from the Muschelkalk (Middle Triassic) of the Germanic Basin—indicators of their ecology, extinctions, and immigrations. *Palaeogeography, Palaeoclimatology, Palaeoecology,* 221(1-2), 7-34.
55. Kucera, M., & Malmgren, B. A. (1998). Differences between evolution of mean form and evolution of new morphotypes: an example from Late Cretaceous planktonic foraminifera. *Paleobiology*, 24(1), 49-63.
56. Landman, N. H., Klofak, S. M., & Sarg, K. B. (2008). Variation in adult size of scaphitid ammonites from the Upper Cretaceous Pierre Shale and Fox Hills Formation (pp. 149-194). Springer Netherlands.
57. Łaska, W., Rodríguez-Tovar, F. J., & Uchman, A. (2017). Evaluating macrobenthic response to the Cretaceous–Palaeogene event: a high-resolution ichnological approach at the Agost section (SE Spain). *Cretaceous Research*, 70, 96-110.
58. Leone, M. F., & Benedetto, J. L. (2019). The Brachiopod *Dalmanella testudinaria* across the End Ordovician Extinction Event in the Cuyania Terrane of Western Argentina. *Ameghiniana*, 56(3), 228-242.
59. Leu, M., Bucher, H., & Goudemand, N. (2019). Clade-dependent size response of conodonts to environmental changes during the late Smithian extinction. *Earth-Science Reviews*, 195, 52-67.
60. Liow, L. H., & Taylor, P. D. (2019). Cope's Rule in a modular organism: Directional evolution without an overarching macroevolutionary trend. *Evolution*, 73(9), 1863-1872.
61. Liu, G., Feng, Q., Shen, J. U. N., Yu, J., He, W., & Algeo, T. J. (2013). Decline of siliceous sponges and spicule miniaturization induced by marine productivity collapse and expanding anoxia during the Permian-Triassic crisis in South China. *Palaios*, 28(8), 664-679.
62. Lockwood, R. (2005). Body size, extinction events, and the early Cenozoic record of veneroid bivalves: a new role for recoveries?. *Paleobiology*, 31(4), 578-590.

63. Luo, G., Lai, X., Shi, G. R., Jiang, H., Yin, H., Xie, S., ... & Wignall, P. B. (2008). Size variation of conodont elements of the *Hindeodus–Isarcicella* clade during the Permian–Triassic transition in South China and its implication for mass extinction. *Palaeogeography, Palaeoclimatology, Palaeoecology,* 264(1-2), 176-187.
64. MacLeod, N., Ortiz, N., Fefferman, N., Clyde, W., Schulter, C., MacLean, J. (2000). Phenotypic response of foraminifera to episodes of global environmental change. In S.J. Culver and P.F. Rawson (Eds), Biotic response to global environmental change: the last 145 Million Years (1st ed., pp. 51-78). Cambridge University Press.
65. Martindale, R. C., & Aberhan, M. (2017). Response of macrobenthic communities to the Toarcian Oceanic Anoxic Event in northeastern Panthalassa (Ya Ha Tinda, Alberta, Canada). *Palaeogeography, Palaeoclimatology, Palaeoecology*, 478, 103-120.
66. Martínez-Díaz, J. L., Phillips, G. E., Nyborg, T., Espinosa, B., de Araújo Távora, V., Centeno-García, E., & Vega, F. J. (2016). Lilliput effect in a retroplumid crab (Crustacea: Decapoda) across the K/Pg boundary. *Journal of South American Earth Sciences*, 69, 11-24.
67. Mattioli, E., & Pittet, B. (2002). Contribution of calcareous nannoplankton to carbonate deposition: a new approach applied to the Lower Jurassic of central Italy. *Marine micropaleontology*, 45(2), 175-190.
68. Mattioli, E., Pittet, B., Petitpierre, L., & Mailliot, S. (2009). Dramatic decrease of pelagic carbonate production by nannoplankton across the Early Toarcian anoxic event (T-OAE). *Global and Planetary Change*, 65(3-4), 134-145.
69. Metcalfe, B., Twitchett, R. J., & Price-Lloyd, N. (2011). Changes in size and growth rate of 'Lilliput'animals in the earliest Triassic. *Palaeogeography, Palaeoclimatology, Palaeoecology,* 308(1-2), 171-180.
70. Monnet, C., Bucher, H., Brayard, A., & Jenks, J. F. (2013). *Globacrochordiceras* gen. nov.(Acrochordiceratidae, late Early Triassic) and its significance for stress-induced evolutionary jumps in ammonoid lineages (cephalopods). *Fossil Record,* 16(2), 197-215.
71. Monnet, C., Bucher, H., Guex, J., Wasmer, M. (2012). Large-scale evolutionary trends of Acrochordiceratidae Arthaber, 1911 (Ammonoidea, Middle Triassic) and Cope's Rule.- *Palaeontology*, 55(1), 87-107.
72. Morten, S. D., & Twitchett, R. J. (2009). Fluctuations in the body size of marine invertebrates through the Pliensbachian-Toarcian extinction event. *Palaeogeography, Palaeoclimatology, Palaeoecology*, 284(1-2), 29-38.
73. Munroe, D.M., Narváez, D.A., Hennen, D., Jacobson, L., Mann, R., Hofmann, E.E., Powell, E.N. and Klinck, J.M. (2016). Fishing and bottom water temperature as drivers of change in maximum shell length in Atlantic surfclams (*Spisula solidissima*). *Estuarine, Coastal and Shelf Science*, 170, 112-122.
74. Murphy, M. A., & Springer, K. B. (1989). Morphometric study of the platform elements of *Amydrotaxis praejohnsoni* n. sp.(Lower Devonian, Conodonts, Nevada). *Journal of Paleontology*, 63(3), 349-355.
75. Mutter, R. J., & Neuman, A. G. (2009). Recovery from the end-Permian extinction event: evidence from "Lilliput Listracanthus". *Palaeogeography, Palaeoclimatology, Palaeoecology*, 284(1-2), 22-28.
76. Nätscher, P. S., Gliwa, J., De Baets, K., Ghaderi, A., & Korn, D. (2023). Exceptions to the temperature–size rule: no Lilliput Effect in end-Permian ostracods (Crustacea) from Aras Valley (northwest Iran). *Palaeontology*, 66(4), e12667.
77. Novack-Gottshall, P. M. (2008). Ecosystem-wide body-size trends in Cambrian–Devonian marine invertebrate lineages. *Paleobiology*, 34(2), 210-228.
78. Nürnberg, S., Aberhan, M., & Krause, R. A. (2012). Evolutionary and ecological patterns in body size, shape, and ornamentation in the Jurassic bivalve *Chlamys (Chlamys) textoria* (Schlotheim, 1820). *Fossil Record*, 15(1), 27-39.
79. O'Dea, A., Hakansson, E., Taylor, P. D., & Okamura, B. (2011). Environmental change prior to the K-T boundary inferred from temporal variation in the morphology of cheilostome bryozoans. *Palaeogeography, Palaeoclimatology, Palaeoecology*, 308(3-4), 502-512.
80. Opazo, L. F., & Twitchett, R. J. (2022). Bivalve body-size distribution through the Late Triassic mass extinction event. *Paleobiology*, 48(3), 420-445.
81. Ortega, L., Celentano, E., Delgado, E., & Defeo, O. (2016). Climate change influences on abundance, individual size and body abnormalities in a sandy beach clam. *Marine Ecology Progress Series*, 545, 203-213.
82. Payne, J. L. (2005). Evolutionary dynamics of gastropod size across the end-Permian extinction and through the Triassic recovery interval. *Paleobiology*, 31(2), 269-290.

83. Payne, J. L., Groves, J. R., Jost, A. B., Nguyen, T., Moffitt, S. E., Hill, T. M., & Skotheim, J. M. (2012). Late Paleozoic fusulinoidean gigantism driven by atmospheric hyperoxia. *Evolution*, 66(9), 2929-2939.
84. Peng, Y., Shi, G. R., Gao, Y., He, W., & Shen, S. (2007). How and why did the Lingulidae (Brachiopoda) not only survive the end-Permian mass extinction but also thrive in its aftermath?. *Palaeogeography, Palaeoclimatology, Palaeoecology*, 252(1-2), 118-131.
85. Piazza, V., Duarte, L. V., Renaudie, J., & Aberhan, M. (2019). Reductions in body size of benthic macroinvertebrates as a precursor of the early Toarcian (Early Jurassic) extinction event in the Lusitanian Basin, Portugal. *Paleobiology*, 45(2), 296-316.
86. Piazza, V., Ullmann, C. V., & Aberhan, M. (2020). Temperature-related body size change of marine benthic macroinvertebrates across the Early Toarcian Anoxic Event. *Scientific reports*, 10(1), 4675.
87. Pietsch, C., Gigliotti, M., Anderson, B. M., & Allmon, W. D. (2023). Patterns and processes in the history of body size in turritelline gastropods, Jurassic to Recent. *Paleobiology*, 49(4), 621-641.
88. Pimiento, C., & Balk, M. A. (2015). Body-size trends of the extinct giant shark *Carcharocles megalodon*: a deep-time perspective on marine apex predators. *Paleobiology*, 41(3), 479-490.
89. Pohle, A., & Klug, C. (2018). Body size of orthoconic cephalopods from the late Silurian and Devonian of the Anti-Atlas (Morocco). *Lethaia*, 51(1), 126-148.
90. Posenato, R., Holmer, L. E., & Prinoth, H. (2014). Adaptive strategies and environmental significance of lingulid brachiopods across the late Permian extinction. *Palaeogeography, Palaeoclimatology, Palaeoecology*, 399, 373-384.
91. Pye, F. (2017). Survival of the smallest? Trends in brachiopod size across the End-Triassic mass extinction. *The Palaeontology Newsletter* 94, 93-96.
92. Qiu, X., Tian, L., Wu, K., Benton, M. J., Sun, D., Yang, H., & Tong, J. (2019). Diverse earliest Triassic ostracod fauna of the non-microbialite-bearing shallow marine carbonates of the Yangou section, South China. *Lethaia*, 52(4), 583-596.
93. Rego, B. L., Wang, S. C., Altiner, D., & Payne, J. L. (2012). Within-and among-genus components of size evolution during mass extinction, recovery, and background intervals: a case study of Late Permian through Late Triassic foraminifera. *Paleobiology,* 38(4), 627-643.
94. Renaud, S., & Girard, C. (1999). Strategies of survival during extreme environmental perturbations: evolution of conodonts in response to the Kellwasser crisis (Upper Devonian). *Palaeogeography, Palaeoclimatology, Palaeoecology*, 146(1-4), 19-32.
95. Reyment, R. A. (1963). Studies on Nigerian upper Cretaceous and lower Tertiary Ostracoda. Part 2. Danian, Paleocene, and Eocene Ostracoda. *Stockholm Contributions in Geology*, 10.
96. Rice, E., Dam, H. G., & Stewart, G. (2015). Impact of climate change on estuarine zooplankton: surface water warming in Long Island Sound is associated with changes in copepod size and community structure. *Estuaries and coasts*, 38, 13-23.
97. Rick, T. C., Reeder-Myers, L. A., Hofman, C. A., Breitburg, D., Lockwood, R., Henkes, G., ... & Ogburn, M. B. (2016). Millennial-scale sustainability of the Chesapeake Bay Native American oyster fishery. *Proceedings of the National Academy of Sciences*, 113(23), 6568-6573.
98. Rita, P., De Baets, K., & Schlott, M. (2018). Rostrum size differences between Toarcian belemnite battlefields. *Fossil Record*, 21(1), 171-182.
99. Rita, P., Nätscher, P., Duarte, L. V., Weis, R., & De Baets, K. (2019). Mechanisms and drivers of belemnite body-size dynamics across the Pliensbachian–Toarcian crisis. *Royal Society open science*, 6(12), 190494.
100. Rogov, M. A. (2020). Infrazonal ammonite biostratigraphy, paleobiogeography and evolution of Volgian craspeditid ammonites. *Paleontological Journal*, 54(10), 1189-1219.
101. Rogov, M. A., Shchepetova, E. V., Ippolitov, A. P., Seltser, V. B., Mironenko, A. A., Pokrovsky, B. G., & Desai, B. G. (2019). Response of cephalopod communities on abrupt environmental changes during the early Aptian OAE1a in the Middle Russian Sea. *Cretaceous Research*, 96, 227-240.
102. Romano, C., Koot, M. B., Kogan, I., Brayard, A., Minikh, A. V., Brinkmann, W., ... & Kriwet, J. (2016). Permian–Triassic Osteichthyes (bony fishes): diversity dynamics and body size evolution. *Biological Reviews*, 91(1), 106-147.
103. Ros-Franch, S., Echevarria, J., Damborenea, S. E., Mancenido, M. O., Jenkyns, H. C., Al-Suwaidi, A., ... & Riccardi, A. C. (2019). Population response during an Oceanic Anoxic Event: The case of Posidonotis (Bivalvia) from the Lower Jurassic of the Neuquen Basin, Argentina. *Palaeogeography, Palaeoclimatology, Palaeoecology*, 525, 57-67.

104. Roy, K., Collins, A. G., Becker, B. J., Begovic, E., & Engle, J. M. (2003). Anthropogenic impacts and historical decline in body size of rocky intertidal gastropods in southern California. *Ecology Letters*, 6(3), 205-211.
105. Salamon, M. A., Brachaniec, T., Brom, K. R., Lach, R., & Trzęsiok, D. (2016). Dwarfism of irregular echinoids (*Echinocorys*) from Poland during the Campanian-Maastrichtian Boundary Event. *Palaeogeography, Palaeoclimatology, Palaeoecology*, 457, 323-329.
106. Salamon, M.A., Brachaniec, T., Paszcza, K., Kolbuk, D., Gorzelak, P. (2023). The role of mass extinction events in shaping the body-size dynamics of fossil crinoids. *Palaeogeography, Palaeoclimatology, Palaeoecology*, 622, 111593.
107. Sallan, L., and Galimberti, A. K. (2015). Body-size reduction in vertebrates following the end-Devonian mass extinction. *Science* 350(6262), 812-815.
108. Sarkar, D., Paul, S., Saha, R., Bardhan, S., Rudra, P. (2022) Body size trends in Trigoniida Bivalves from the Mesozoic Kutch, India. *Palaios*, 37, 89-103.
109. Schaal, E. K., Clapham, M. E., Rego, B. L., Wang, S. C., & Payne, J. L. (2016). Comparative size evolution of marine clades from the Late Permian through Middle Triassic. *Paleobiology*, 42(1), 127-142.
110. Schmidt, D.N., Thomas, E., Authier, E., Saunders, D., Ridgwell, A. (2018). Strategies in times of crisis - insights into the benthic foraminiferal record of the Palaeocene-Eocene Thermal Maximum. *Phil. Trans. R. Soc*., 376, 20170328.
111. Schmidt, D. N., Thierstein, H. R., Bollmann, J., & Schiebel, R. (2004). Abiotic forcing of plankton evolution in the Cenozoic. *Science*, 303(5655), 207-210.
112. Song, H., J. Tong & Z. Q. Chen (2011). Evolutionary dynamics of the Permian-Triassic foraminifer size: Evidence for Lilliput effect in the end-Permian mass extinction and its aftermath. *Palaeogeography, Palaeoclimatology, Palaeoecology*, 308, 98-110.
113. Shukla, S.K., Romero, O.E. (2018). Glacial valve size variation of the Southern Ocean diatom *Fragilariopsis kerguelensis* preserved in the Benguela Upwelling System, southeastern Atlantic. *Palaeogeography, Palaeoclimatology, Palaeoecology*, 499, 112-122.
114. Sigurdsen A., Hammer O. (2016). Body size trends in the Ordovician to earliest Silurian of the Oslo Region. *Palaeogeography, Palaeoclimatology, Palaeoecology,* 443, 49-56.
115. Sogot, C. E., Harper, E. M., & Taylor, P. D. (2014). The Lilliput effect in colonial organisms: cheilostome bryozoans at the Cretaceous–Paleogene mass extinction. Plos One, 9(2), e87048.
116. Takahashi, A. (2005). Responses of inoceramid bivalves to environmental disturbances across the Cenomanian/Turonian boundary in the Yezo forearc basin, Hokkaido, Japan. *Cretaceous Research*, 26(4), 567-580.
117. Tewfik, A., Babcock, E. A., Appeldoorn, R. S., & Gibson, J. (2019). Declining size of adults and juvenile harvest threatens sustainability of a tropical gastropod, *Lobatus gigas*, fishery. Aquatic Conservation: *Marine and Freshwater Ecosystems*, 29(10), 1587-1607.
118. Trubovitz, S., & Stigall, A. L. (2018). Ecological revolution of Oklahoma's rhynchonelliform brachiopod fauna during the Great Ordovician Biodiversification Event. *Lethaia*, 51(2), 277-285.
119. Wade, B. S., & Olsson, R. K. (2009). Investigation of pre-extinction dwarfing in Cenozoic planktonic foraminifera. *Palaeogeography, Palaeoclimatology, Palaeoecology*, 284(1-2), 39-46.
120. Wei, F. (2019). Conch size evolution of Silurian–Devonian tentaculitoids. *Lethaia*, 52(4), 454-463.
121. Wiest, L. A., Buynevich, I. V., Grandstaff, D. E., Terry Jr, D. O., Maza, Z. A., & Lacovara, K. J. (2016). Ichnological evidence for endobenthic response to the K–Pg event, New Jersey, USA. *Palaios*, 31(5), 231-241.
122. Wiest, L. A., Buynevich, I. V., Grandstaff, D. E., Terry Jr, D. O., & Maza, Z. A. (2015). Trace fossil evidence suggests widespread dwarfism in response to the end-Cretaceous mass extinction: Braggs, Alabama and Brazos River, Texas. *Palaeogeography, palaeoclimatology, palaeoecology*, 417, 105-111.
123. Wilson-Brodie, R. J., MacLean, M. A., & Fenberg, P. B. (2017). Historical shell size reduction of the dogwhelk (*Nucella lapillus*) across the southern UK. *Marine biology*, 164(9), 190.
124. Witts, J. D., Landman, N. H., Hopkins, M. J., & Myers, C. E. (2020). Evolutionary stasis, ecophenotypy and environmental controls on ammonite morphology in the Late Cretaceous (Maastrichtian) Western Interior Seaway, USA. *Palaeontology*, 63(5), 791-806.

125. Wu, K., Tian, L., Liang, L., Metcalfe, I., Chu, D., & Tong, J. (2019). Recurrent biotic rebounds during the Early Triassic: biostratigraphy and temporal size variation of conodonts from the Nanpanjiang Basin, South China. *Journal of the Geological Society*, 176(6), 1232-1246.
126. Yamaguchi, T., Norris, R. D., & Bornemann, A. (2012). Dwarfing of ostracodes during the Paleocene–Eocene Thermal Maximum at DSDP Site 401 (Bay of Biscay, North Atlantic) and its implication for changes in organic carbon cycle in deep-sea benthic ecosystem. *Palaeogeography, Palaeoclimatology, Palaeoecology*, 346, 130-144.
127. Yang, L., Dai, X., Liu, X., Feng, Y., Jiang, S., Wang, F., Song, H., Tian, L., Song, H. (2024) Foraminiferal Extinction and Size Reduction during the Permian-Triassic Transition in Southern Tibet. *Journal of Earth Science*, 35(6), 1799-1809.
128. Zhang, X., Li, S., Song, Y., Gong, Y. (2020). Size reduction of conodonts indicates high ecological stress during the late Frasnian under greenhouse climate conditions in South China. *Palaeogeography, Palaeoclimatology, Palaeoecology,* 556, 109909.
129. Zhang, Y., Shi, G. R., He, W. H., Wu, H. T., Lei, Y., Zhang, K. X., ... & Xiao, Y. F. (2016). Significant pre-mass extinction animal body-size changes: evidences from the Permian–Triassic boundary brachiopod faunas of South China. *Palaeogeography, Palaeoclimatology, Palaeoecology*, 448, 85-95.
130. Zhang, Z. T., Sun, Y. D., Wignall, P. B., Fu, J. L., Li, H. X., Wang, M. Y., & Lai, X. L. (2018). Conodont size reduction and diversity losses during the Carnian Humid Episode in SW China. *Journal of the Geological Society*, 175(6), 1027-1031.

**SI Appendix References II (References included from the Hunt et al. (2015) dataset)**

1. Baumfalk, Y. A., Troelstra, S. R., Ganssen, G., & Van Zanen, M. J. (1987). Phenotypic variation of *Globorotalia scitula* (Foraminiferida) as a response to Pleistocene climatic fluctuations. *Marine geology*, 75(1-4), 231-240.
2. Biolzi, M. (1991). Morphometric analyses of the Late Neogene planktonic foraminiferal lineage *Neogloboquadrina dutertrei*. *Marine Micropaleontology*, 18(1-2), 129-142.
3. Blaj, T., J. Hendricks, J. R. Young, and E. Rehnberg. (2010). The Oligocene nannolith Sphenolithus evolutionary lineage: morphometrical insights from the palaeo-equatorial Pacific Ocean. *Journal of Micropalaeontology,* 29, 17-35.
4. Bornemann, A., & Mutterlose, J. (2006). Size analyses of the coccolith species *Biscutum constans* and *Watznaueria barnesiae* from the Late Albian “Niveau Breistroffer”(SE France): taxonomic and palaeoecological implications. *Geobios*, 39(5), 599-615.
5. Bralower, T. J., & Parrow, M. (1996). Morphometrics of the Paleocene coccolith genera *Cruciplacolithus*, *Chiasmolithus*, and *Sullivania*: a complex evolutionary history. *Paleobiology*, 22(3), 352-385.
6. Cheetham, A. H. (1968). Morphology and systematics of the bryozoan genus *Metrarabdotos*. *Smithsonian Miscellaneous Collections*, 153,121
7. Christensen, W. K. (2000). Gradualistic evolution in *Belemnitella* from the middle Campanian of Lower Saxony, NW Germany. *Bulletin of the Geological Society of Denmark,* 47, 135-163.
8. Drooger, C. W., and D. S. N. Raju. (1978). Early Miogypsinoides in Kutch, western India (I). *Proceedings of the Koninklijke Nederlandse van Wetenschappen series B*, 81, 186-201.
9. Dzik, J. (2008). Evolution of morphogenesis in 360-million-year-old conodont chordates calibrated in days. *Evolution & Development*, 10, 769-777.
10. Fermont, W. J. J. (1982). Discocyclinidae from Ein Avedat (Israel). *Utrecht Micropaleontological Bulletins,* 27, 1-152.
11. Geary, D. H., G. Hunt, I. Magyar, and H. Schreiber. (2010). The paradox of gradualism: phyletic evolution in two lineages of lymnocardiid bivalves (Lake Pannon, central Europe). *Paleobiology*, 36, 592-614.
12. Hayami, I. (1984). Natural history and evolution of Cryptopecten (a Cenozoic-Recent pectinid genus). *Bulletin of the University Museum, University of Tokyo*, 24, 149.
13. Hodell, D. A., & Vayavananda, A. (1993). Middle Miocene paleoceanography of the western equatorial Pacific (DSDP site 289) and the evolution of Globorotalia (Fohsella). *Marine Micropaleontology*, 22(4), 279-310.
14. Hunt, G. (2007). The relative importance of directional change, random walks, and stasis in the evolution of fossil lineages. *Proceedings of the National Academy of Sciences*, 104(47), 18404-18408.

15. Jones, D. (2009). Directional evolution in the conodont *Pterospathodus*. *Paleobiology*, 35, 413-431.
16. Kaim, A. (2002). Gradual evolution of the Early Cretaceous marine gastropod Rissoina lineage in central Poland. *Acta Palaeontologica Polonica*, 47, 667-672.
17. Kelley, P. H. (1983). Evolutionary patterns of eight Chesapeake group molluscs: evidence for the model of punctuated equilibria. *Journal of Paleontology,* 57, 581-598.
18. Kellogg, D. E. (1975). The role of phyletic change in the evolution of *Pseudocubus vema* (Radiolaria). *Paleobiology*, 1, 359-370.
19. Kellogg, D. E. (1980). Character displacement and phyletic change in the evolution of the radiolarian subfamily Artiscinae. *Micropaleontology*, 26, 196-210.
20. Kim, K., H. D. Sheets, and C. E. Mitchell. (2009). Geographic and stratigraphic change in the morphology of *Triarthrus beckii* (Green) (Trilobita): A test of the Plus ca change model of evolution. *Lethaia,* 42, 108-125.
21. Knappertsbusch, M. (2000). Morphologic evolution of the coccolithophorid *Calcidiscus leptoporus* from the Early Miocene to Recent. *Journal of Paleontology*, 74(4), 712-730.
22. Knappertsbusch, M. (2007). Morphological variability of *Globorotalia menardii* (planktonic foraminifera) in two DSDP cores from the Caribbean Sea and the Eastern Equatorial Pacific. *Carnets de Géologie/Notebooks on Geology*, (A04), 1-34.
23. Kucera, M., & Kennett, J. P. (2002). Causes and consequences of a middle Pleistocene origin of the modern planktonic foraminifer *Neogloboquadrina pachyderma* sinistral. *Geology*, 30(6), 539-542.
24. Laagland, H. (1990). Cycloclypeus in the Mediterranean Oligocene (Doctoral dissertation, Utrecht University).
25. Lazarus, D., R. Scherer, P., and D. R. Prothero. (1985). Evolution of the radiolarian species-complex Pterocanium: a preliminary survey. *Journal of Paleontology,* 59, 183-220.
26. Malmgren, B., Kucera, M., Ekman, G. (1996). Evolutionary Changes in Supplementary Apertural Characteristics of the Late Neogene. *PALAIOS*, 11, 192-206.
27. Malmgren, B. A., Berggren, W. A., & Lohmann, G. P. (1983). Evidence for punctuated gradualism in the Late Neogene *Globorotalia tumida* lineage of planktonic foraminifera. *Paleobiology*, 9(4), 377-389.
28. Malmgren, B. A., & Kennett, J. P. (1981). Phyletic gradualism in a Late Cenozoic planktonic foraminiferal lineage; DSDP Site 284, southwest Pacific. *Paleobiology*, 7(2), 230-240.
29. Motoyama, I. (1997). Origin and evolution of *Cycladophora davisiana* Ehrenberg (radiolaria) in DSDP Site 192, Northwest Pacific. *Marine Micropaleontology*, 30(1-3), 45-63.
30. Pearson, P. N., & Ezard, T. H. (2014). Evolution and speciation in the Eocene planktonic foraminifer *Turborotalia*. *Paleobiology*, 40(1), 130-143.
31. Pokorny, V. (1966). La variation de la taille moyenne chez les Ostracodes comme indice paleoecologique. *Ecologae Geologicae Helvetiae* 59, 269-276.
32. Renaud, S., & Schmidt, D. N. (2003). Habitat tracking as a response of the planktic foraminifer *Globorotalia truncatulinoides* to environmental fluctuations during the last 140 kyr. *Marine Micropaleontology*, 49(1-2), 97-122.
33. Schultz, M. G. (1979). Morphometrisch-variationsstatistische Untersuchungen zur Phylogenie der Belemniten-Gattung *Belemnella* im Untermaastricht NW-Europas. *Geologisches Jahrbuch*, A 47:3-157.
34. Sorhannus, U. (1990). Punctuated morphological change in a Neogene diatom lineage:“local” evolution or migration?. *Historical Biology*, 3(4), 241-247.
35. Sorhannus, U., Fenster, E. J., Burckle, L. H., & Hoffman, A. (1988). Cladogenetic and anagenetic changes in the morphology of *Rhizosolenia praebergonii* Mukhina. *Historical Biology*, 1(3), 185-205.
36. Springer, K. B., & Murphy, M. A. (1994). Punctuated stasis and collateral evolution in the Devonian lineage of *Monograptus hercynicus*. *Lethaia*, 27(2), 119-128.
37. Tanabe, K. (1973). Evolution and mode of life of *Inoceramus (Sphenoceramus) naumanni* Yokoyama emend., an Upper Cretaceous bivalve. *Transactions and Proceedings of the Palaeontological Society of Japan, new series,* 92, 163-184.
38. Tanabe, K. (1977). Functional evolution of *Otoscaphites puerculus* (Jimbo) and *Scaphites planus* (Yabe), Upper Cretaceous ammonites. *Memoirs of the Faculty of Science Kyushu University Series D Geology,* 23, 367-407.
39. Theriot, E. C., Fritz, S. C., Whitlock, C., & Conley, D. J. (2006). Late Quaternary rapid morphological evolution of an endemic diatom in Yellowstone Lake, Wyoming. *Paleobiology*, 32(1), 38-54.

40. Tiraboschi, D., & Erba, E. (2010). Calcareous nannofossil biostratigraphy (Upper Bajocian–Lower Bathonian) of the Ravin du Bès section (Bas Auran, Subalpine Basin, SE France): evolutionary trends of *Watznaueria barnesiae* and new findings of “Rucinolithus” morphotypes. *Geobios*, 43(1), 59-76.
41. Wildenborg, A. F. B. (1991). Evolutionary aspects of the Miogypsinids in the Oligo-Miocene carbonates near Mineo (Sicily). *Utrecht Micropaleontological Bulletins*, (41).
